# Multiscale mechanistic modelling of the host defense in invasive aspergillosis reveals leukocyte activation and iron acquisition as drivers of infection outcome

**DOI:** 10.1101/2021.06.08.447590

**Authors:** Henrique AL Ribeiro, Luis Sordo Vieira, Yogesh Scindia, Bandita Adhikari, Matthew Wheeler, Adam Knapp, William Schroeder, Borna Mehrad, Reinhard Laubenbacher

**Author notes:** Co-last authors (Borna Mehrad, Reinhard Laubenbacher).

## Abstract

Aspergillus species are ubiquitous environmental molds, the spores of which are inhaled daily by most humans. Immunocompromised hosts can develop an invasive infection resulting in high mortality. There is, therefore, a pressing need for host-centric therapeutics for this infection. To address this need, we created a multi-scale computational model of the infection, focused on its interaction with the innate immune system and iron, a critical nutrient for the pathogen. The model, parameterized using published data, was found to recapitulate a wide range of biological features and was experimentally validated *in vivo*. Conidial swelling was identified as critical in fungal strains with high growth, whereas the siderophore secretion rate is an essential prerequisite for establishment of the infection in low-growth strains. In immunocompetent hosts, high growth, high swelling probability, and impaired leukocyte activation lead to a high conidial germination rate. Similarly, in neutropenic mice, high fungal growth was achieved through synergy between high growth rate, high swelling probability, slow leukocyte activation, and high siderophore secretion. In summary, the model reveals a small set of parameters related to fungal growth, iron acquisition, and leukocyte activation as key determinants of the fate of the infection.

## Introduction

Invasive aspergillosis is a human infection with increasing incidence, related to the use of immunosuppressive therapies, such as cancer chemotherapy and immunosuppression medications [1]. More recently, it has also been observed that 10% to 14% of critically ill patients with COVID-19 developed invasive aspergillosis [2, 3]. Mortality remains high, 30-60% in recent surveys [4], despite advances in diagnostics and therapy. Increasing triazole resistance in this infection [5] has raised the specter of a “perfect storm” [6] in an increasing population of susceptible individuals with a diminished repertoire of treatment options.

The research presented here was motivated by the search for host-centric interventions in immuno-compromised patients that can be used in combination with antifungal treatments. An important mechanism in innate immunity is the sequestration of iron from pathogens, a nutrient critical for nearly all organisms. A well-established literature supports the concept that the “battle over iron” is characteristic of the host’s attempt to attenuate microbial growth during many infections [7]. Iron is particularly relevant to the pathogenesis of aspergillosis [8]. The iron sequestration feature of the innate immune response involves several intertwined processes that unfold across spatial and temporal scales. This makes it challenging to assess the effect of perturbations of individual mechanisms on infection dynamics. A computational model that captures the key mechanisms, broadly reflects the underlying immune biology, and is well-validated, can play an essential role in hypothesis generation and the discovery of emergent properties of the immune response.

Several models related to respiratory *Aspergillus* infections and their pathology have been previously published. For example, agent-based models have shown the necessity of chemotactic signals for proper fungal clearance [9, 10]. Our own work includes a model of the innate immune response to *A. fumigatus*, showing that a key determinant of infection is the range at which macrophages can detect the fungus [11], and an intracellular regulatory network linking iron metabolism to oxidative stress in a fungal cell [12]. The model presented here is parameterized entirely with information from the literature, rather than through data fitting, and is validated by showing that it can recapitulate a wide range of experimental data and features reported in the literature that were not used in its construction, as well as experimental data generated for the purpose of model validation. The model was then used to identify major drivers of the growth of fungal burden, providing potential targets for intervention.

## Materials and methods

### A computational model of the immune response to invasive aspergillosis

The model is an agent-based model of invasive pulmonary aspergillosis scaled to a mouse lung, the experimental system used in this study, focusing on the “battle over iron” between host and fungus. It integrates the critical players in the early immune response and the known mechanisms that govern their behavior and interactions. It is divided into six conceptual components: space and time, molecules, host cells, *Aspergillus fumigatus*, interactions between different cells, and cells and molecules, and iron metabolism, as briefly described here.

### Space and time

A three-dimensional space representing a small portion of a mouse lung is divided into a discrete grid of one thousand voxels (10 voxels in each of 3 dimensions), representing a total volume of 6.4 × 10^−2^ *μ*L. Each voxel has an edge length of 40 *μm* (6.4 × 10^−5^*μL*). Cells and molecules have no space coordinates other than the voxel in which they are located at a given time. This approach is similar to that used in the general immune modeling platform C-IMMSIM [13]. The space has periodic boundary conditions, and simulated time progresses in discrete steps of two minutes.

### Molecules

The model includes five kinds of molecules: cytokines (IL6, IL10, TGF, TNF, CXCL2, CCL4), a siderophore (TAFC), iron carrier molecules (transferrin and lactoferrin), iron, and the hormone hepcidin. The cytokines are subject to a half-life of one hour [14–20]. Furthermore, all the molecules are subject to a constant exchange between the simulated volume and the serum (system). Iron in this model is used only as a temporary buffer for the transference of iron between dying cells (i.e., macrophages, *A. fumigatus*) and iron-carrier molecules.

The concentration of a molecule in one voxel is called the local concentration, and we will refer to the concentration across the whole simulated space as the global concentration. In contrast, the serum concentration (i.e., outside the simulator) is the systemic concentration. Equation 1 determines the flux of molecules between the serum and the simulated (local) space. For the cytokines lactoferrin and TAFC, we assume as a simplification that the systemic concentration *x*_*system*_ is zero. Therefore, these molecules are constantly flowing in the direction of the serum, increasing their decay rate. The systemic level of hepcidin and transferrin is dynamically calculated as follows:

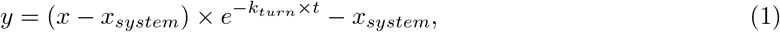

where *x*_*system*_ is the molecule’s systemic concentration (see terminology above), *x* is the local concentration, *k*_*turn*_ is the turnover rate, and *t* is the time-step length (2 min).

Equations 2 and 3 compute the systemic levels of hepcidin and transferrin. Note that the result of the first equation (2) feeds into the second one (3). However, the first equation needs the systemic levels of IL6 as input. However, in this simulator, we only have the local and global levels of IL6. According to Goncalves, SM *et al*. 2017 [21], a reasonable estimate of the systemic level of IL6 is one-half of its global level. The equations are:

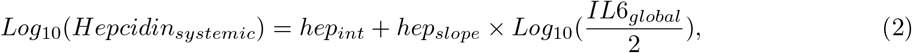

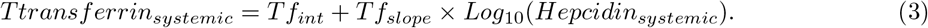

These equations are based on data from Tabbah S *et al*. 2018 [22], correlating systemic levels of IL-6 to systemic levels of hepcidin, and Moran-Lev H *et al*. 2018 [23] correlating transferrin and hepcidin. To ensure biologically meaningful values we only evaluate these equations if *IL*6_*global*_ *>* 1.37 × 10^−10^*M*, and *Hepcidin*_*systemic*_ *>* 10^−8^*M*.

All these molecules, except iron, diffuse through space, modeled using the Alternating Direction Implicit (ADI) method with a periodic boundary condition [24]. The rationale for periodic boundary conditions is that the simulation covers a small area amid a large infected area. Therefore, the concentration of molecules across the boundaries should be similar.

### Host Cells

There are three types of host cells: type II pneumocytes, macrophages, and neutrophils. The host cells can assume several different states and transition from one to another upon interacting with other agents or molecules. Figure 1 shows the graph of host cell states. We introduce the states “activating” and “inactivating” to model the delay between signal and phenotype change. For a review of macrophage activation/inactivation, see Duque, GA & Descoteaux, A 2014 [25] and Gordon, S 2003 [26].

**Fig 1.**
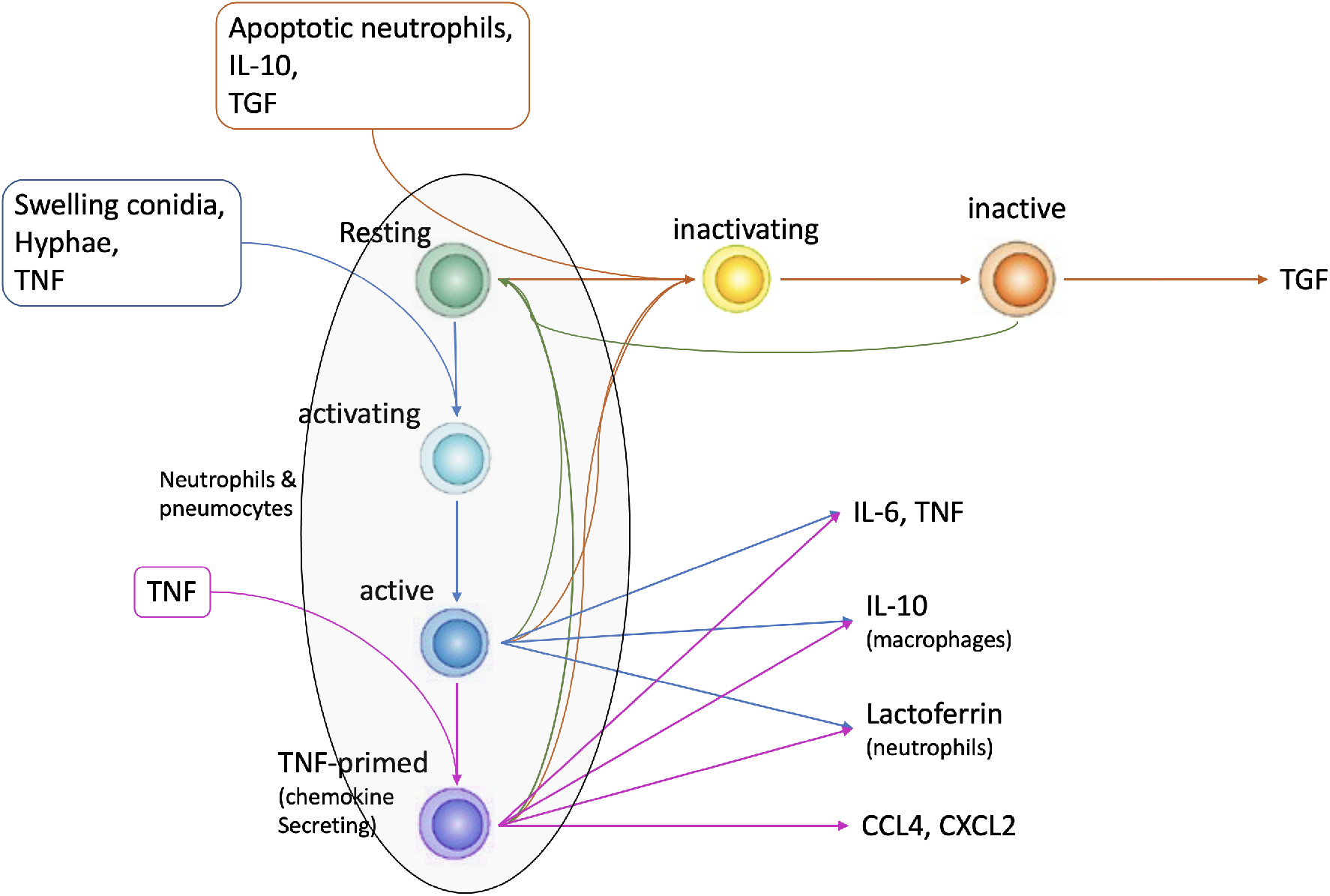
Figure showing host cells state changes. The paragon host cells in the model are the macrophages. Therefore, this figure represents the whole state space of a macrophage. The other cells (neutrophils and type II pneumocytes) have a subset of the states macrophages have (see the area in the ellipses). By default, host cells are in a resting state. Swelling conidia, hyphae, or TNF causes them to transition to an activating (intermediate) state and then to the active state. Active host cells secrete TNF, IL-6, and IL-10. Extra priming with TNF makes host cells secrete chemokines as well (CCL4 and CXCL2). All macrophages return to a resting state after 6 hours (180 iterations) in the absence of a continuous stimulus. Apoptotic neutrophils, IL-10, or TGF-*β*, cause macrophages (including activated macrophages) to transition to an anti-inflammatory TGF-*β*-secreting state. Active macrophages (blue and purple) can kill hyphae whereas resting and anti-inflammatory ones cannot.

Leukocytes are free to move through the space and can be recruited as well. Recruitment is done according to Equation 4:

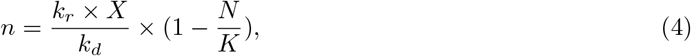

where *N* is the current number of cells in the simulator, K is the carrying capacity, *k*_*r*_ is the global recruitment rate, *k*_*d*_ is the dissociation constant of the chemokine, *X* is the global amount of the chemokine (*i*.*e*., the average concentration of chemokine in the simulator), and *n* is the average number of cells to be recruited. This number is used by a Poisson random number generator to decide how many cells will be recruited. Macrophages and neutrophils have half-lives of 24 and 6 hours, respectively [27, 28]. The quantity of cells in the simulator is a balance between the number of cells recruited according to Equation 4 and the number of cells that die. Macrophages are recruited by CCL4 and neutrophils by CXCL2.

In the absence of chemokines, cells move randomly, while in their presence, they tend to move to the voxels with higher chemokine concentrations. The rate of movement is constant, and the cells will, on average, traverse a fixed number of voxels per time step. In the presence of chemokines, each voxel receives a weight according to Equation 5:

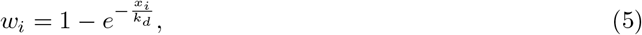

where *x*_*i*_ is the chemokine concentration in neighboring voxel *i, w*_*i*_ is the corresponding weight of this voxel, and *k*_*d*_ is the chemokine dissociation constant. The cell will then move to a neighboring voxel (*v*_*i*_) with probability proportional to the voxel weight (*p*_*i*_ *∝ w*_*i*_).

### Aspergillus

In the model, *Aspergillus fumigatus* has three life stages: resting conidia, swelling conidia, and hyphae. The hyphae are more or less continuous structures divided by septae [29]. Each of these subdivisions is a multinucleated cell-like structure, referred to as a hyphal cell for simplicity.

In previous work, a dynamic gene regulatory network of iron uptake by *Aspergillus fumigatus* was developed [12], in the form of a Boolean network, that is used here as a component model, with minor adjustments. While the original network had a TAFC node activating the node LIP, the labile iron pool, representing metabolically available iron, we modified it as follows. Instead, when TAFC is activated, the cell secretes TAFC. Later on, if the cell is expressing the siderophore receptors, the cell takes up TAFCBI (TAFC Bound to Iron) from the environment. The TAFCBI uptake increases the cell’s total iron pool. The LIP node then becomes a function of the iron pool. Using Equation 6, we activate LIP if the iron pool is high. Resting conidia do not produce or take up TAFC in our model. We update the Boolean network every 30 minutes (every 15 iterations of the tissue-scale model) [30].

In simulations, *Aspergillus fumigatus* starts out as a pool of resting conidia; after 4 hours these start swelling with a half-life of 6 hours (see Table S1) - that is, half the conidia swell after 6h. Beyond that, it takes 2 hours until they become able to grow into hyphal cells. However, growth is controlled by LIP. That is, if LIP is off, hyphae cannot develop, consistent with the known importance of iron for hyphal growth.

Although hyphal growth is a continuous process, the model uses a discrete approximation. A tip cell can produce another tip cell (elongation), while a sub-tip cell can form a 45-degree branch (subapical branch) [29, 31] with a 25% probability. Other cells cannot originate new cells unless their neighbors are killed, and they become tip or sub-tip cells again.

### Interactions

Table S2 displays the interactions between all cell types and molecular species. Note that cells/agents may interact with each other in more than one way, depending on their state. For example, the agent “Afumigatus” (that represents *Aspergillus fumigatus*) has the states “resting conidia,” “swelling conidia,” and “hyphae.” The interaction between this agent and a neutrophil will be different depending on this state (see Table S2). Due to software engineering considerations, we consider the secretion of molecules a kind of interaction between a cell and a molecule. Therefore, interactions between molecules and cells can also be of two kinds: receptor binding (interaction in the biological sense) and secretion. Note that, in Table S2, macrophages need to be active in order to interact with hyphae, while neutrophils do not. For agents to interact with each other, they must be in physical proximity, *i*.*e*., in the same voxel. Interactions between cells and molecules follow Equation 6:

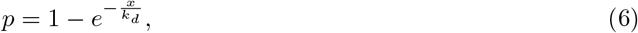

where *x* is the molecule concentration, *k*_*d*_ is its dissociation constant, and *p* is the probability of receptor activation. Likewise, interactions between molecules (i.e., reactions) follow the Michaelian equation (Equation 7):

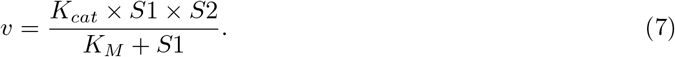

### Scaling from the simulated space to the whole mouse lung

A pair of mouse lungs is assumed to have a volume of 1mL [32], containing 2.3 × 10^5^ macrophages in the alveolar lumen [33] and 1 × 10^7^ type-II alveolar epithelial cells [34]. The simulated space is 6.4 × 10^−2^*μL*, thus containing 15 macrophages and 640 type-II epithelial cells initially. A high dose inoculum (10^7^) is used for initialization. However, according to Pritchard, JN *et al*. 1985 [35], inoculated material distributes unevenly, with ≈ 1*/*3 of the lung infected and the remainder clear. Scaling appropriately, the simulated space should have 1920 conidia. To scale neutrophils, one can also use the fact that infection is limited to ≈ 1*/*3 of the lung. In other words, to convert the number of neutrophils and fungal spores in the simulated space to the whole lung number, one needs to multiply by 5028.

## Experimental methods

### Neutrophil depletion and induction of aspergillosis

All experiments were performed in accordance with the National Institutes of Health and Institutional Animal Care and Use Guidelines and were approved by the Animal Care and Use Committee of the University of Florida. Eight week-old male and female C57Bl/6 mice were purchased from the Jackson Laboratory and housed under specific pathogen-free conditions in the animal facilities of the University of Florida, and infected with Aspergillus as previously described by us [36]. Briefly, neutrophils were transiently depleted with an intraperitoneal injection of 400*μg* of anti-Ly6G antibody (clone 1A8, BioXcell) in 0.5ml saline. A cohort of mice received an equivalent amount of isotype control antibody (rat IgG2a, Clone 2A3, BioXcell), a day prior to intratracheal inoculation with 7 × 10^6^ *Aspergillus* conidia.

### Flow Cytometry

Mouse lung flow cytometry was performed as described in [37]. Briefly, lungs were digested in a mixture of 200 *μg/mL* DNaseI and 25 *μg/mL* Liberase TM for 30 mins at 370 °*C*. The digested lungs were serially passed through 70 and 40 *μm* filters to collect the single-cell suspension. After red blood cell lysis, cells were counted, and 1.5 × 10^6^ cells were stained with a fixable APC Cy-7 conjugated live dead stain (Thermo Fisher) in PBS for 20 mins. After washing with FAC buffer, cells were incubated with anti-CD16/32 (Fc block, clone 93; eBioscience, San Diego, CA) and stained with PerCP-conjugated anti-CD45 (30-F11), FITC-conjugated anti-CD11b (M1/70), PE-conjugated CD64 (X54-5/7.1), PECy7-conjugated anti-CD11c (N418), V450-conjugated anti-MHCII (I-A/I-E), APC-conjugated anti-CD24 (M1/69), BV605-conjugated anti-Ly6g (1A8), BV711-conjugated Ly6c (HK 1.4), Texas Red-conjugated Siglec F (E50-2440). Flow cytometry data were acquired using 14 color BD Fortessa (BD Biosciences, San Jose, CA). 500,000 events /samples were acquired and analyzed with FlowJo software 9.0 (Tree Star Inc., Ashland, OR).

### Bronchoalveolar lavage fluid cytokine measurement

BALF IL-6 and CXCL2 levels were measured using commercial ELISA kits (Invitrogen), as per manufacturers’ instructions.

## Results

### Model validation

This model was completely parameterized with data from the literature (see Supplementary Materials), thereby assuring a much broader validity than could be obtained through parameter fitting to a small collection of experimental time-course measurements. We have validated the model in two ways. Firstly, we show that it provides a good qualitative fit with several time-course data sets in the literature that were not used in model calibration. We used a collection of papers that report time-series of critical variables present in the model, such as neutrophils, TNF, IL6, and colony-forming units (CFU). These values are compared with those predicted by model simulation. These data are used to test if the model can reproduce the reported levels of the different variables and, most importantly, their timing. None of the papers selected for validation were used to calibrate the model.

Figure 2 exhibits the comparison of simulation results with literature data. One can observe that the simulator correctly captures the timing of cell counts and cytokine levels. Furthermore, CFU dynamics (Figure 2B) is also accurately captured by the model. Concerning the exact levels of these cells and cytokines, the mismatches between our simulator and the data in Figure 2 are within the range of variation between the different experiments reported in the literature (Table 1). Figure 3 shows that our model correctly predicts the fall in CXCL2 upon injection of anti-TNF-*α*.

**Table 1.** Table showing extended literature measurements of *Aspergillus fumigatus* outcome parameters. All the papers in this table report data in BAL upon 24 hours post-infection and inoculate mice with ≈ 10^7^ conidia. Column 1 shows the reference; column 2 reported measurements of TNF; column 3 IL-6; column 4 neutrophils; and column 5 *log*_10_ of CFU.

| Reference | TNF | IL-6 | Neutrophils | $\log_{10}(CFU)$ |
| --- | --- | --- | --- | --- |
| Bhatia, S *et al.* 2011 [38] | | | $9.60 \pm 0.14 \times 10^5$ | |
| Brieland, JK *et al.* 2001 [39] | $3027 \pm 194pg/mL$ | | | $5.56 \pm 0.10$ |
| Cenci, E *et al.* 2001 [43] | $1602 \pm 297pg/mL$ | $348 \pm 52pg/mL$ | | |
| Dubourdeau, M *et al.* [44] | $923 \pm 174pg/mL$ | $64 \pm 18pg/mL$ | | |
| Doung, M *et al.* 1998 [40] | | $460 \pm 8pg/mL$ | | $6.24 \pm 0.16$ |
| Gresnigt, MS *et al.* 2016 [45] | | $364 \pm 47pg/mL$ | $5.42 \pm 1.64 \times 10^5$ | |
| Hohl, TM *et al.* 2005 [46] | | | $2.30 \pm 0.92 \times 10^6$ | |
| Teschner, D *et al.* 2019 [47] | $592 \pm 48pg/mL$ | $1964 \pm 313$ | $4.04 \pm 1.25 \times 10^5$ | $4.38 \pm 0.38$ |
| **Average ± std-dev** | **$1536 \pm 1079$ pg/mL** | **$676 \pm 748$ pg/mL** | **$1.05 \pm 0.87 \times 10^6$** | **$5.39 \pm 0.94$** |

**Fig 2.**
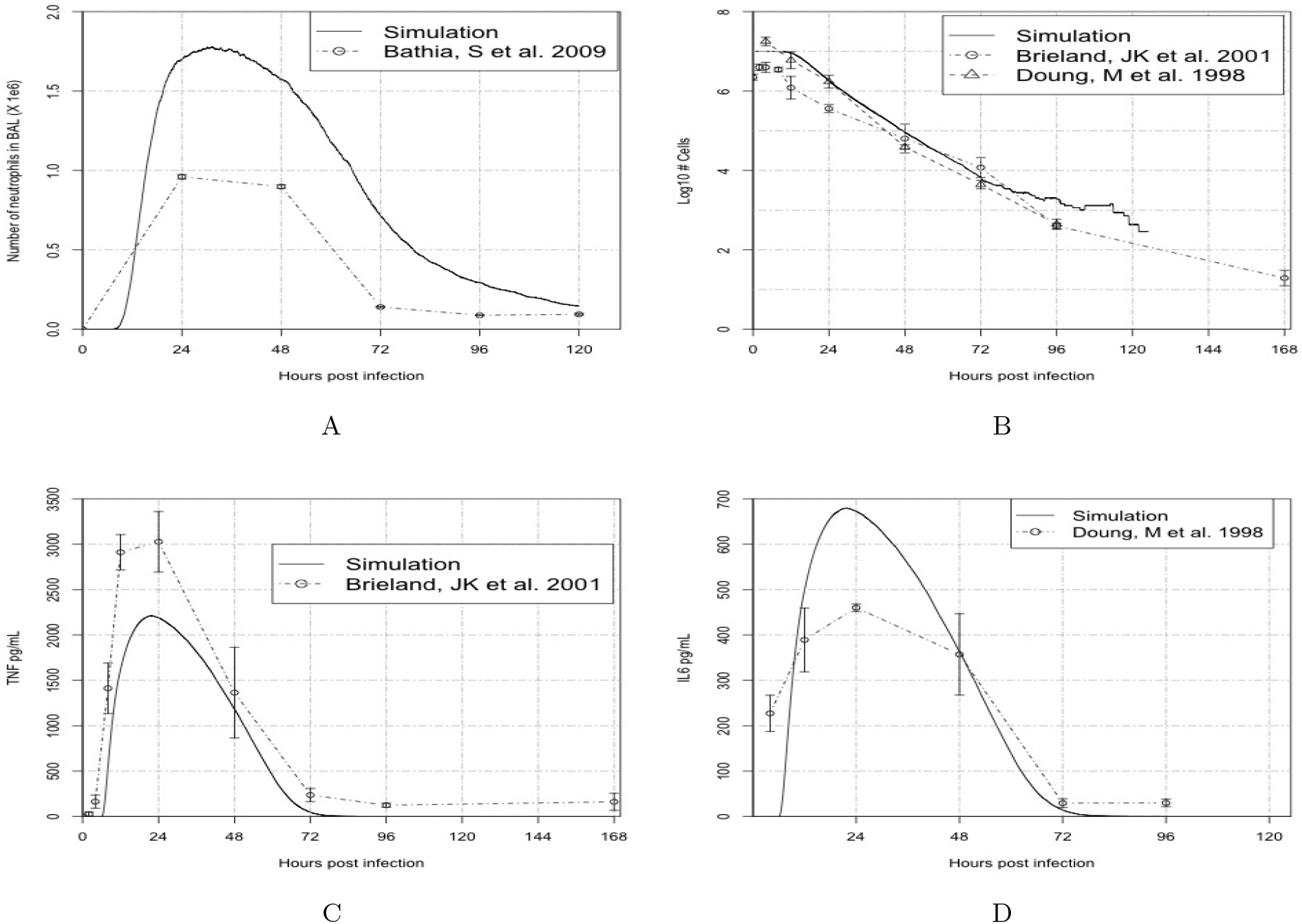
Figure showing the comparison of simulated data with data reported in the literature. To produce this figure, 36 simulations were performed, starting with an average of 1920 conidia, 15 macrophages, and 640 epithelial cells. Figure 2A: simulated time series of neutrophils and a time series reported by Bhatia, S *et al*. 2011 [38]. Figure 2B: simulated time series of conidia and time series reported by Brieland, JK et al. 2001 [39] and Doung, M *et al*. 1998 [40]. Figure 2C: simulated time series of TNF and time series reported by Brieland, JK *et al*. 2001 [39]. Figure 2D: simulated time series of IL-6 and time series reported by Doung, M *et al*. 1998 [40].

**Fig 3.**
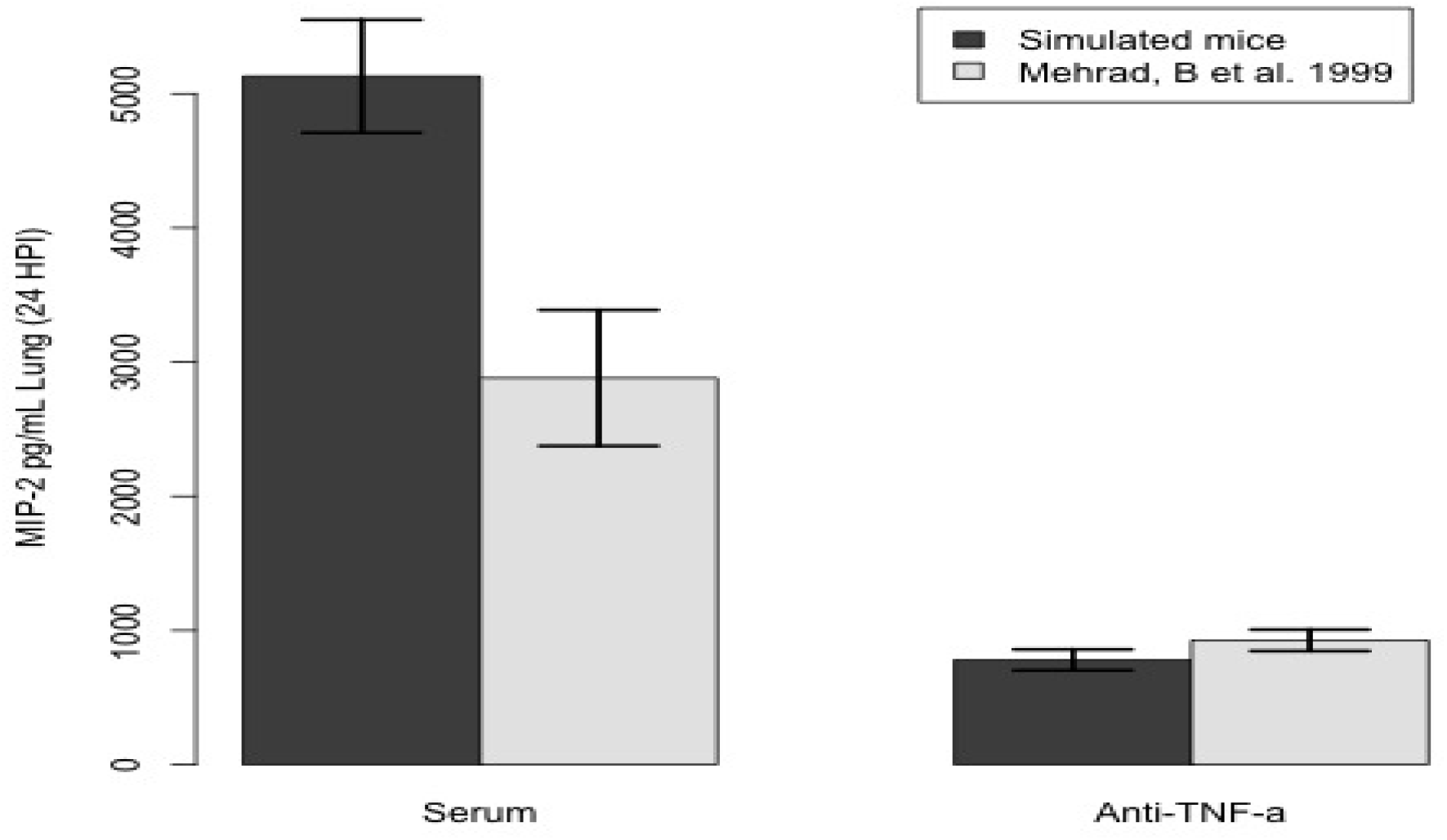
Figure showing the comparison of simulated data with data reported by Mehrad, B *et al*. 1999 [41]. Mice were injected with serum or antibody (anti-TNF) concentration of 2 × 10^−8^ M, reaction rate 1.72 × 10^8^ *M * s*^−1^ (Kcat/Km), and half-life of 5 days, 24 before infection. To produce this figure, 36 simulations were performed, initialized with an average of 1920 conidia, 15 macrophages, and 640 epithelial cells.

Next, we compared model simulations with a published experiment of mice injected with anti-TNF-*α*. This cytokine is one of the critical drivers of the immune response and one of the key molecules inducing the secretion of chemokines. In Mehrad et al. 1999 [41], the level of chemokines fell 24 hours post-infection upon injection with an anti-TNF antibody. To reproduce this experiment, we estimated the affinity of an antibody for a protein antigen with data from the literature [42]. Note that those estimates are for a generic protein antigen and not for TNF specifically.

We did an extensive literature search for data concerning *Aspergillus fumigatus* infection in mice. We compiled this data in Table 1. It represents BAL measurements of IL-6, TNF, CFU, and neutrophils 24 hours post-infection with 10^7^ conidia. One can observe that these data have significant variability. To understand how our simulated data compare to these measurements, we sampled parameters with Latin Hypercube Sampling (LHS) and ran 1200 simulations. Figure 4 displays the comparison between several literature measurements, the simulator with the default parameters (Table S1), and the simulator with LHS parameters. The predictions made by our simulator are within the range of variation between data reported in the literature. Note that the simulator can also reproduce the variability in biological data. This variability is due to differences in hosts, different fungal strains, and experimental conditions. These conditions sometimes generate different but qualitatively equivalent outcomes. Our simulator is robust to a wide range of parameters, with variability similar to that observed in the literature Figure; see 4.

**Fig 4.**
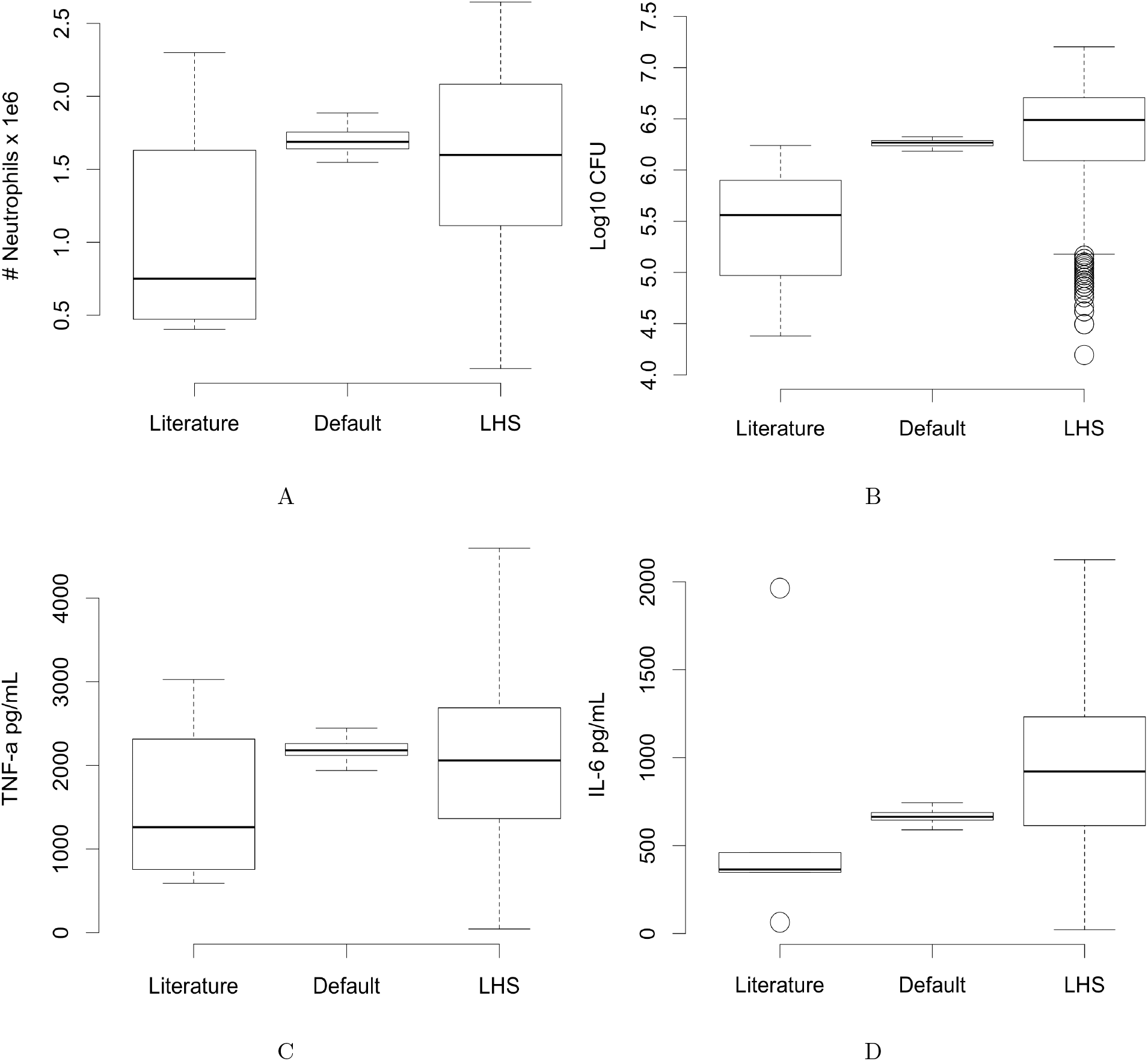
Figures showing the comparison of simulated data with extended literature reported data (Table 1). Literature: data from the literature (Table 1). Default: simulator run with default values. LHS simulator run with parameters sampled with LHS. To generate the “Default,” we run 36 simulations with the default parameter values of the simulator; see Supplementary Table S1. To generate the “LHS,” we run 1200 simulations with LHS sampled around the default values.

### *In vivo* validation of simulation results

As discussed above, due to the high variability in experimental design (fungal dose, techniques of infection, etc.) and in the measurements summarized in Table 1, we decided to further validate the model results, in particular the ability of the simulator to reproduce temporal dynamics, with an *in vivo* experimental design that most closely resembles the simulator setup. We infected immunocompetent mice with 7 × 10^6^ conidia intratracheally and measured cytokines and leukocytes from 0-72 hours post-infection (see description in the Experimental Methods section). Figure 5 shows that our simulator can correctly predict the timing of the immune response, as indicated by levels of IL-6, neutrophils, macrophages, and CXCL2.

**Fig 5.**
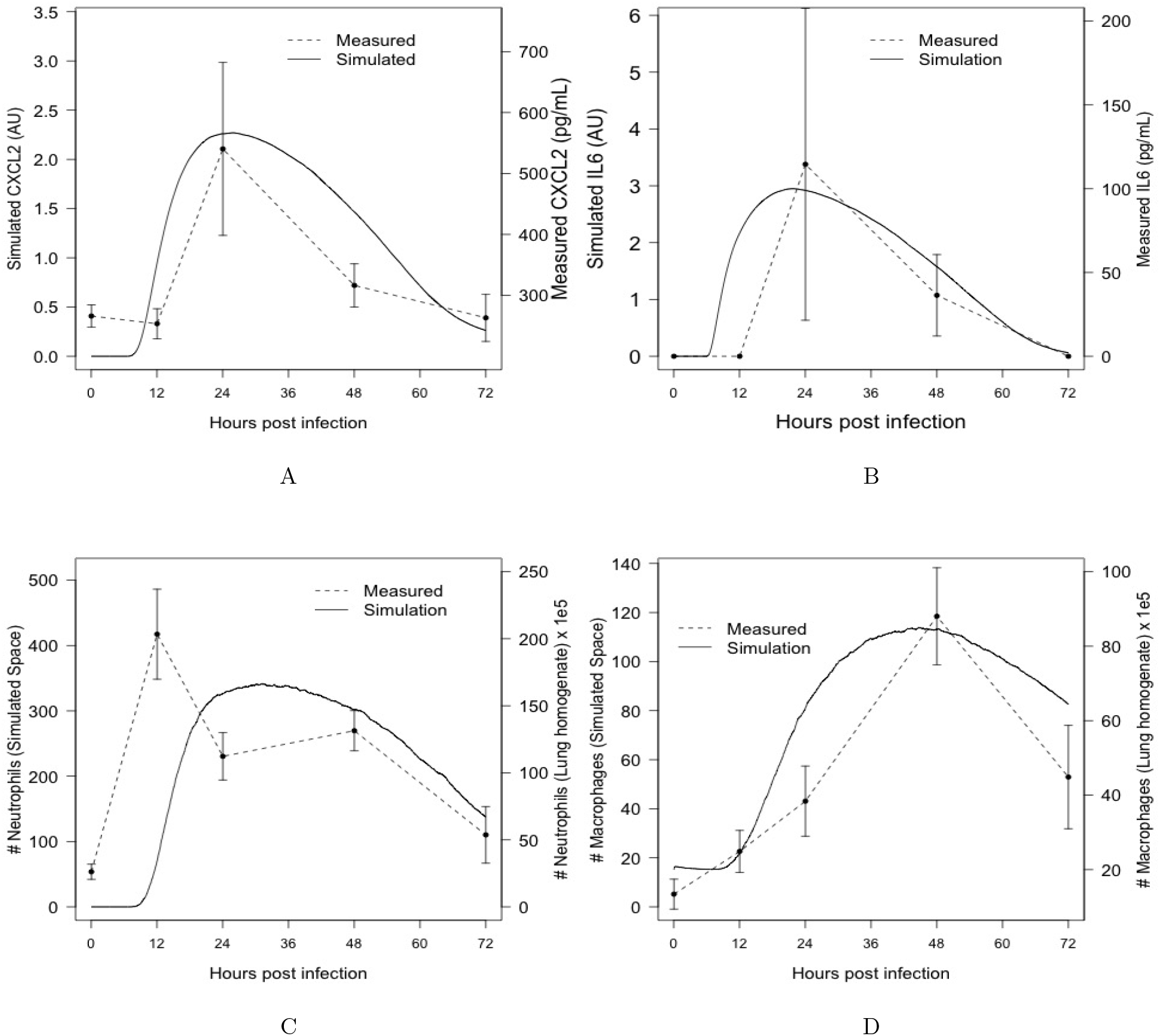
Figure showing the comparison of our experimental data on immunocompetent mice with simulation results. For this purpose, 36 simulations were performed, starting with an average of 1920 conidia, 15 macrophages, and 640 epithelial cells. Figure 5A: Comparison of simulated time series of CXCL2 with experimental data measured in BAL. Figure 5B: Comparison of simulated time series of IL-6 with experimental data measured in BAL. Figure 5C: Comparison of the number of neutrophils in the simulated space with the number of neutrophils in lung homogenate. Figure 5D: Comparison of the number of macrophages/monocytes in simulated space with the number of monocytes in lung homogenate. Experimental data refer to mice infected with 7 × 10^6^ conidia.

Figure 5 shows remarkable agreement between the timing of cytokines and cells measured by our own experiments and the dynamics produced by the simulator. To recall, ABMs are not continuous-time models, since they advance in discrete time steps. The actual time these steps correspond to is estimated by considering the events that take place from one time step to the next. In our case, we estimated that our time steps correspond to two minutes of simulated time. Thus, the agreement in timing is a key step in validating model predictions, since the timing of events in the immune response to this infection is crucial for infection outcome and determination of interventions.

### Identification of drivers of fungal burden

In order to demonstrate the usefulness of the model as a discovery tool for new immunological insights, we use it to investigate the significance of the different mechanisms for fungal growth over time, by measuring their effect on the maximal fungal burden encountered over a 24 hour time span. For this purpose, we conducted the following simulation experiment. Using Latin Hypercube Sampling, as before, we chose 1200 parameter sets. For each of these, we carried out simulation runs with both immunocompetent and neutropenic hosts and counted the number of live hyphal cells over a 24 hour time period. The maximum number encountered during this simulation run was reported as “fungal burden.” (Note that after initial uninhibited growth, the immune response may cause a subsequent drop in the number of live hyphal cells.)

We carried out a classification of parameters and their influence on fungal growth rate, encoded by a collection of classification trees in Figures S2 and S3 in the Supplementary Materials. The results are summarized in Table 2.

**Table 2.**
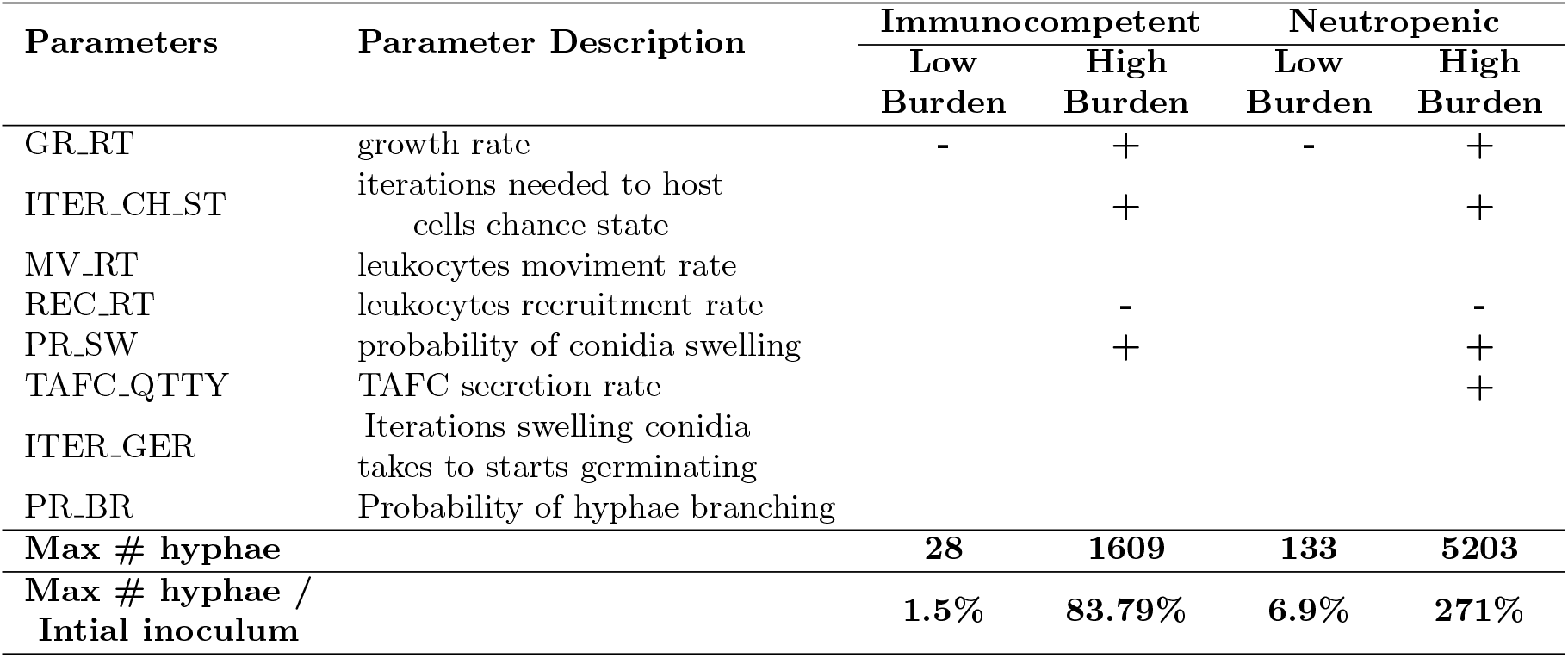
Table summarizing the classification results from Figures S2 and S3. The table shows qualitatively the groups of parameters that lead to the extremes of high and low germination and hyphal proliferation over 24 hours of simulated time. Intermediate conditions are not presented here. The plus and minus signs represent, qualitatively, the classifications. For example, in column 4 (immunocompetent, high burden), the three plus signs and one minus sign indicate that the corresponding partition tree (Figures S2) partitioned the data set by the fungal growth rate, activation rate, swelling rate, and recruitment rate, and selected the upper part of the first three partitions (plus sign) and the lower part of the last (minus sign). In a qualitative sense, this means that that class (immunocompetent, high burden) is associated with high growth rate, activation rate, and swelling rate, and with low recruitment rate. The partitioning hierarchy is not represented in this table, as it is meant to be simply a general summary. Some of the parameters shown in the table do not contribute to the extreme cases reported here; they are kept for completeness. These parameters, however, play a role in differentiating intermediate cases of fungal burden (Figures S2 and S3).

| Parameters | Parameter Description | Immunocompetent |  | Neutropenic |  |
| --- | --- | --- | --- | --- | --- |
|  |  | Low Burden | High Burden | Low Burden | High Burden |
| GR_RT | growth rate | - | + | - | + |
| ITER_CH_ST | iterations needed to host cells chance state |  | + |  | + |
| MV_RT | leukocytes moviment rate |  |  |  |  |
| REC_RT | leukocytes recruitment rate |  | - |  | - |
| PR_SW | probability of conidia swelling |  | + |  | + |
| TAFC_QTTY | TAFC secretion rate |  |  |  | + |
| ITER_GER | Iterations swelling conidia takes to starts germinating |  |  |  |  |
| PR_BR | Probability of hyphae branching |  |  |  |  |
| **Max # hyphae** |  | **28** | **1609** | **133** | **5203** |
| **Max # hyphae / Intial inoculum** |  | **1.5%** | **83.79%** | **6.9%** | **271%** |

Our analysis shows that eight model parameters are most strongly correlated to fungal burden. As would be expected, the most critical parameter is the intrinsic growth propensity of a given fungal strain to grow, (GR RT), that remains fixed for a given fungal strain [48]. Fixing this parameter, we then asked which other parameters were associated with high, respectively low, fungal burden over a 24-hour period. To do this, we measured the variation of the square of the correlation *r*^2^ between the model parameters and fungal burden as the growth rate increases. Figure 6 shows the variation of *r*^2^ for the seven most important parameters (not including intrinsic growth rate) indicated by our analysis in Table 2.

**Fig 6.**
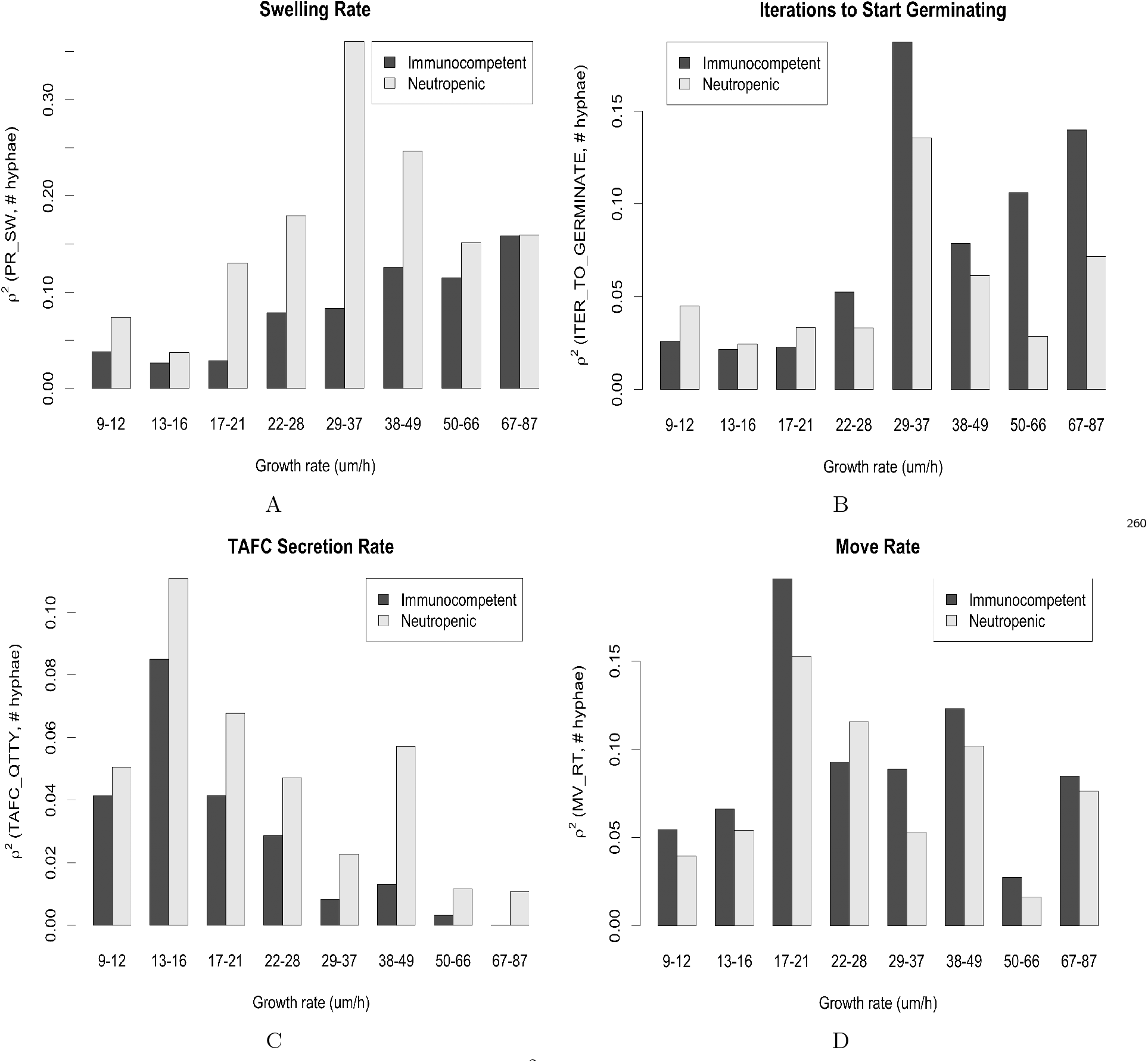
Variation of the square of the correlation *r*^2^ between the critical model parameters and fungal burden, with the variation of the growth rate. We started with a dataset of 1200 samples generated with LHS and then partitioned this dataset by growth rate. The x-axis shows the range of the growth rate in each partition. We then calculate the correlation between parameters and fungal burden in each partition. Fungal burden was measured as the maximum number of hyphal cells measured over the course of 24 hours of simulation. A: Variation of *r*^2^ between the swelling rate and fungal burden. B: Variation of *r*^2^ between germination rate and fungal burden. C: Variation of *r*^2^ between the TAFC secretion rate and fungal burden. D: Variation of *r*^2^ between the leukocyte movement rate and fungal burden.

Figure 6 shows that, as the intrinsic growth rate of the fungus increases, the relative importance of the seven other parameters changes. As an example, Figure 6D shows that when the fungal growth rate is very high or very low, the leukocyte movement rate is weakly correlated with fungal burden. However, this parameter is critical in determining the actual fungal burden in strains with an intrinsic growth rate near the default 29 *μm/h* (Table S1). The leukocyte movement rate is an intrinsic feature of the immune system. However, according to Figure 6D, this feature is more or less critical, depending on the virulence of the strain, being most important for a moderately virulent strain (21 −28*μm/h*). This might explain some of the variance in the course of the infection between hosts.

Figure 6 shows that overall the seven parameters tend to be more critical when the intrinsic growth rate is close to its default value. That is, their importance wanes as the growth rate becomes very high or very low. However, the importance of the swelling rate, measured as the average number of time steps needed for conidia to germinate, and the average time to immune cell activation have a bias towards a high growth rate (Figure 6A, B, and D). On the other hand, the TAFC secretion rate has a bias towards a low growth rate (Figure 6C). Finally, the monocyte recruitment rate is not sensitive to intrinsic growth rate variation (Figure S4C).

The results in Figure 6 also show that activation, germination, and recruitment rates have a bigger influence in immunocompetent hosts (Figure 6B, D, and Figure S4C). Meanwhile, swelling rate, TAFC secretion rate, and branching probability are more crucial in neutropenic hosts (Figure 6A, C, and Figure S4A). Finally, the leucocyte movement rate is approximately equally important in neutropenic and immunocompetent hosts (Figure 6D).

## Discussion

Understanding the innate immune response to pathogens is of the utmost importance for designing effective therapeutic interventions. With the increasing resistance of pathogens to anti-microbial drugs, it is imperative to explore host-centric therapeutics. This is the motivation for the work presented here. The goal was to understand some of the primary components of the innate immune response to fungal pathogens. In order to limit the immense complexity of mechanisms involved, the computational model presented here is focused on an essential component of nutritional immunity, the “battle over iron” between the host and the fungus in the context of a respiratory infection. The component of the immune response considered here involves many players, ranging from immune and fungal cells to molecular species such as cytokines, iron, and chemokines. It integrates events at the intracellular, tissue, organ, and system levels and is governed by several intertwined feedback loops that create complex dynamics.

Without a computational model that captures relevant biology and is parameterized in a way that makes it more broadly valid and credible, it would be challenging to understand the interplay between the different components and make predictions about the effect of various perturbations. This paper describes a model that satisfies these criteria and can serve as the basis for future investigations. It is one of the most comprehensive models of this infection, parameterized entirely with information from the literature, and is validated using experimental data specifically generated for this purpose. It is also shown that it is broadly valid by verifying that it reproduces a wide range of experimental data reported in the literature (and not used for model calibration). This approach is different from the commonly used method of fitting the model parameters to one or more time courses of experimental data.

As an important application of the model to understanding key features of respiratory fungal infections, we used it to study the importance of model parameters to fungal burden, an important metric for disease progression. The model predicts that different characteristics of the immune system take on different significance in controlling fungal burden, depending on the virulence of the fungal strain in question. However, other factors can also significantly influence fungal burden, such as the siderophore secretion rate and certain aspects of the immune response. These factors also have different degrees of importance, depending on the intrinsic growth rate of the fungal strain.

It was pointed out in Schrettl M et al. 2004 [49] that TAFC is a critical virulence factor. Likewise, in subsequent work, Schrettl M et al. 2007 [50] show that an *Aspergillus* knockout strain without intra- and extracellular siderophores has completely attenuated virulence. Figure 6C shows that the TAFC secretion rate is more important when the growth rate is low. In this case, fewer cells are secreting this siderophore. Consequently, the TAFC concentration becomes a bottleneck. In fact, our analysis shows that the concentration of TAFC bound to iron in neutropenic hosts with a high growth rate is 60% higher than in simulations with a low growth rate (result not shown). For immunocompetent hosts, a smaller and less significant difference was observed.

Interestingly, the classification trees (Figure S3, Table 2) show that a high TAFC secretion rate is also necessary for achieving high fungal burden. However, in this case, the TAFC secretion rate acts only in synergy with other parameters. Inhibiting TAFC secretion alone, nevertheless, can break the synergy and reduce fungal burden.

Conidial swelling is the first step to germination and subsequent hyphal growth. However, this is also the time when the fungus become visible to the immune system. Therefore, there are two competing forces upon swelling. The fact that swelling is a necessary step before growth is advantageous for the fungus. This is evidenced by the positive correlation between the swelling rate and fungal burden (results not shown). However, the fact that swelling makes the conidia visible to the immune system and that conidia are easier to kill than hyphae creates a disadvantage for the fungus. This is evidenced by the bias towards a high growth rate (Figure 6A) in this case. That means that swelling is more advantageous if, upon swelling, hyphae develop quickly. A recent model published by Ewald, J *el al*. 2021 [51] came to similar conclusions.

Interestingly, the parameter ITER_GER that controls the time conidia take to germinate after swelling - with two hours being the default value [50] - has a negative correlation with growth rate (result not shown). This negative correlation reinforces that conidia become susceptible to the immune system and need to germinate rapidly upon swelling.

The ability of leucocytes to locate fungal cells is crucial for controlling the infection [9, 11]. The factors that affect it are chemotaxis and movement rate. Figure 6D shows that the leukocyte movement rate is one of the most prominent parameters to control fungal burden with *r*^2^ around 0.2 when the growth rate is in the range 21-28 *μm/h*. However, our analysis also indicates saturation. When the growth rate is too high, the immune system is overwhelmed by fungal growth. Conversely, if the growth rate is too low, the immune system quickly gains the upper hand.

## Availability of code and data

The model code and all experimental data used for model validation are available in the Github repository https://github.com/NutritionalLungImmunity/IPA_model

## Supporting information

**S1 File**. Contains supplementary information on many aspects of the model, model parameters, and simulation.

## Acknowledgments

This work was supported by the following grants: NIH 1U01EB024501-01, NSF CBET-1750183, and NIH 1 R01AI135128-01. R.L. was also partially supported by NIH Grant 1R01GM127909-01, B.M. by American Heart Association Grant 18TPA34170486, and B.A. by NIH Grant NIH T90-DE021989.

## Supplementary Material

## S1 The Parameters

**Table S1:**
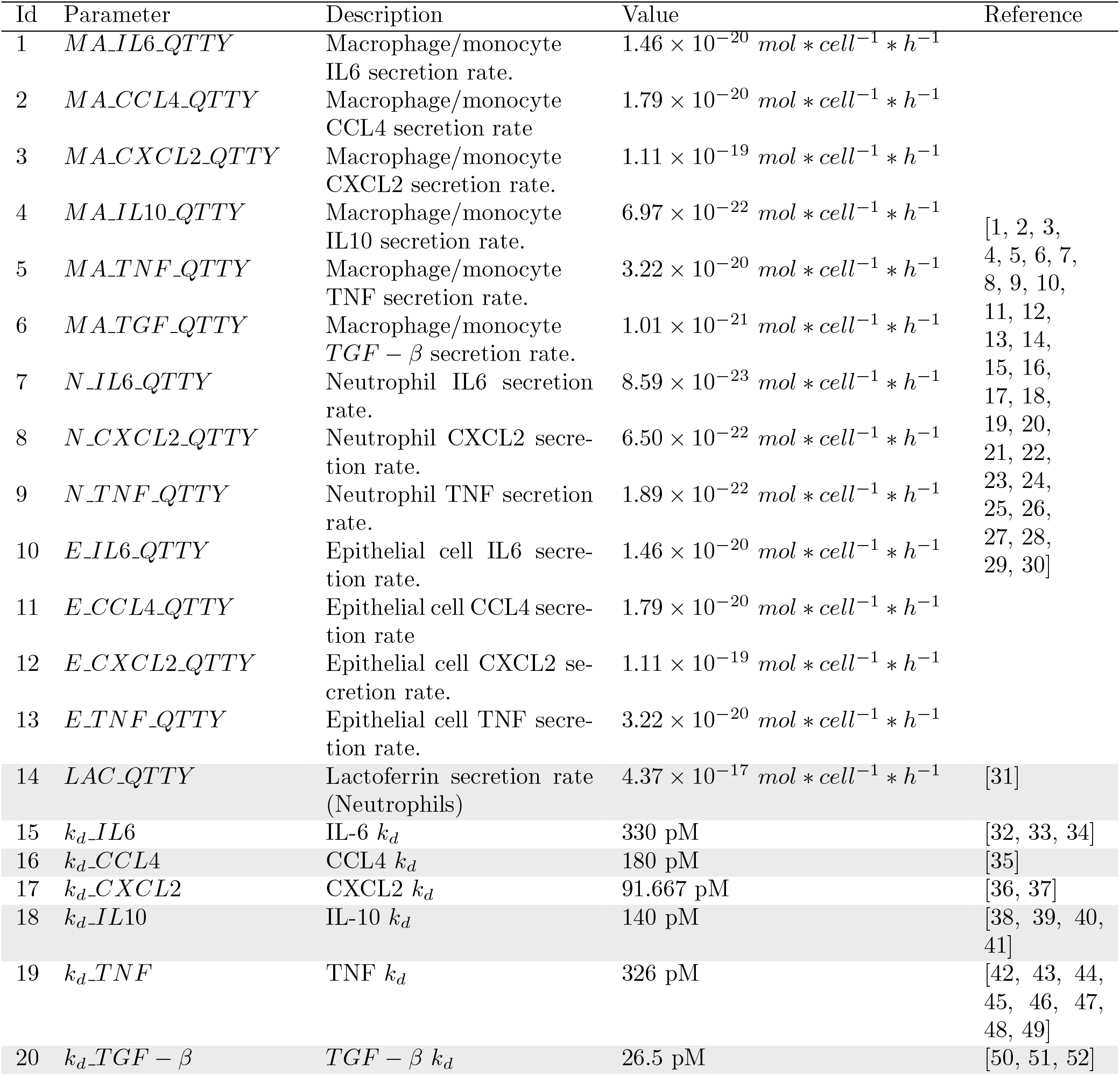
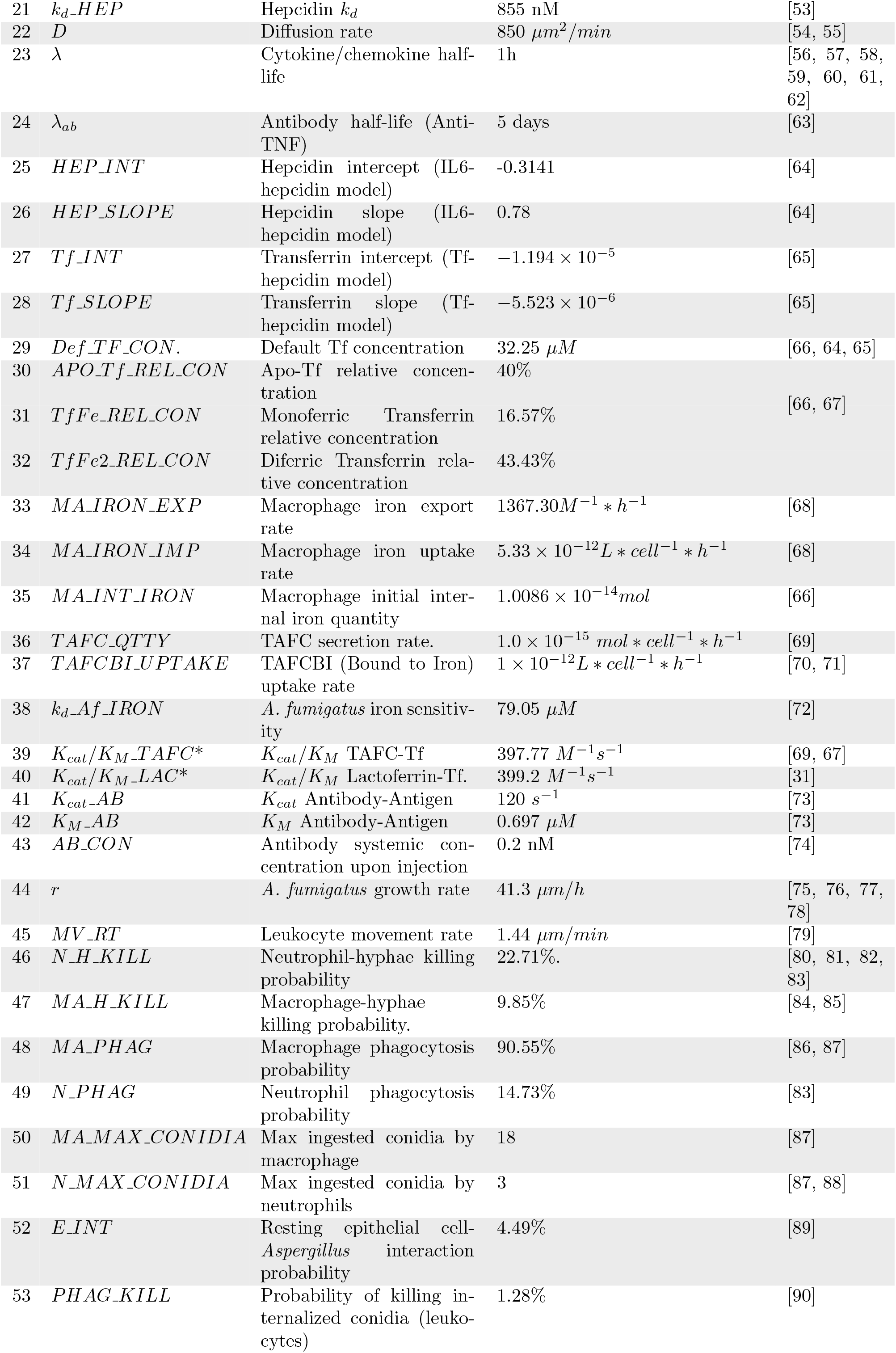
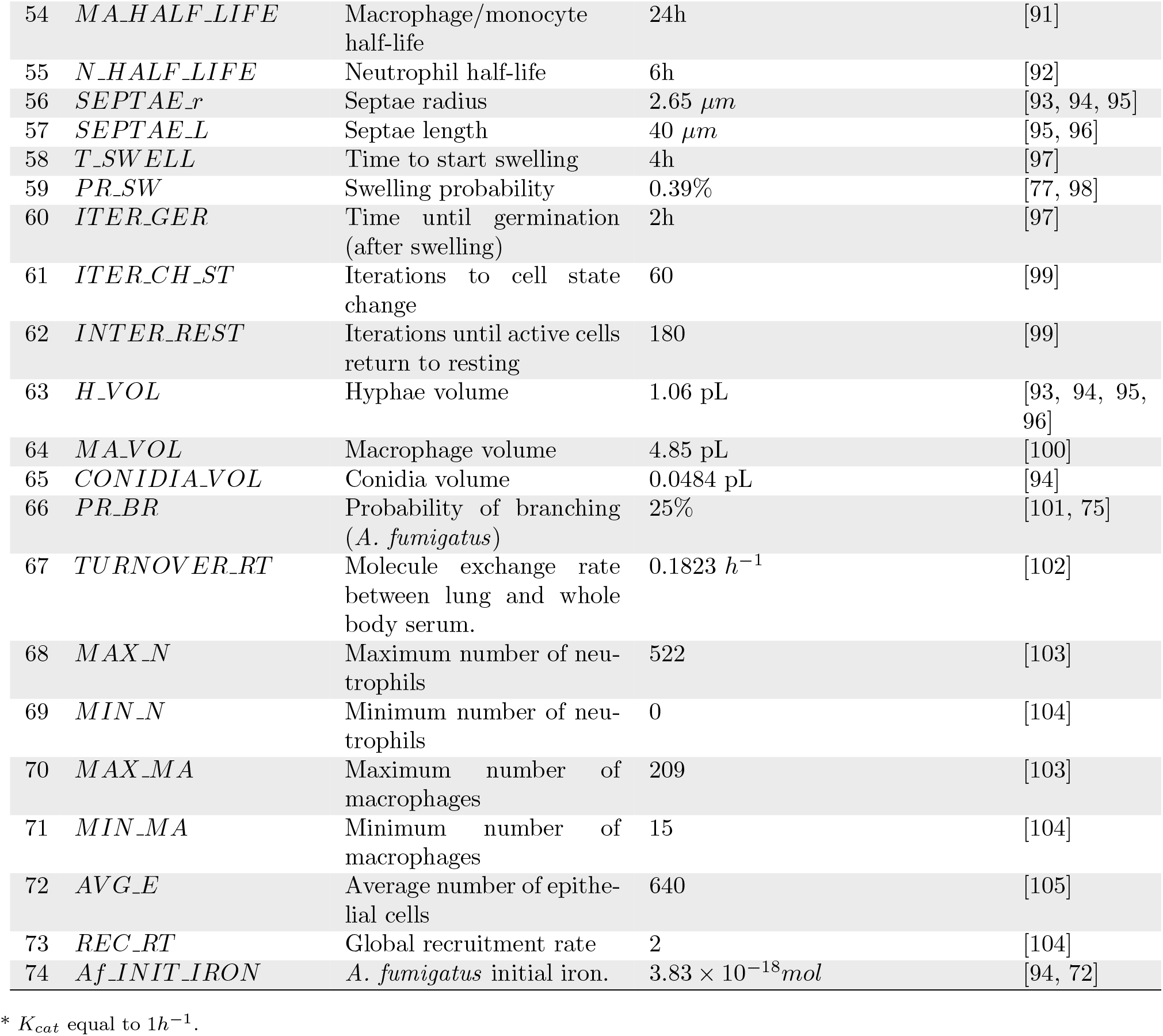
Table with the model parameters. Parameters 1-14 were obtained in a unified manner as described in the manuscript. As mentioned in the manuscript, “macrophage” should be interpreted as “macrophage/monocyte.” Probabilities of phagocytosis, killing, and interaction refer to the likelihood of an event succeeding in one iteration if the appropriate conditions apply. The maximum number of cells and the average number of epithelial cells is computed over the whole simulated space.

## S2 Parameter Acquisition

The model parameters are described in Table S1. In some cases, acquiring these values involved modeling, simplification, and some assumptions. In some cases, we use the MATLAB App Grabit. With this App, one can extract values directly from graphs and pictures. In cases, where more than one measurement is available, the value reported is the median.

### S2.1 Cytokine and chemokine secretion rate

The selection of these rates was done using a collection of papers that report the secretion of cytokines in response to *β* − *glucan, A. fumigatus*, and, in some cases, LPS as a positive control. Each of these papers reports levels of two or more cytokines after monocyte or macrophage exposure with the respective stimulus. Because only papers that reported at least two cytokines were used, it was possible to construct a relationship of relative secretion rates. For instance, note that across experimental procedures, the level of IL-6 is approximately 45% of the level of TNF, while the level of IL-10 is approximately 4.7% of the level of IL-6, etc. Therefore, this relationship specifies that if we secrete *y* of TNF, then 0.45 × *y* IL6 will be secreted.

This procedure was adopted because not all papers, notably those with *β*-glucan, can be quantified. In other words, a response against *β*-glucan is qualitatively similar to a response against live *A. fumigatus*. However, it is not known which concentration of *β*-glucan corresponds to which dose of fungus. With this procedure, however, one can use any piece of data with the implication that one has to fit this relationship to the actual secretion rate. However, having the actual secretion rate for only a few or even one cytokine is enough.

For this purpose, some papers for neutrophils and epithelial cells were used that compare cytokine secretion in these cells with macrophages. Therefore, in the model, these cells are scaled versions of macrophages. Neutrophils secrete 5.9% of what macrophages secret, and epithelial cells secrete the same as macrophages. However, none of these cells secrete IL-10 and TGF-*β*.

The lactoferrin parameter was acquired independently, and it is a rough approximation based on the amount of the protein a neutrophil carries.

### S2.2 TAFC secretion and uptake rates

To calculate the TAFC secretion rate, Equation S2.1 was used to model the experiment of Hissen, AHT *et al*. 2004 [69]. This equation is a surrogate model for our simulator. It models conidia swelling and then secreting TAFC:

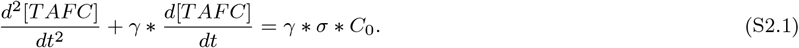

In Equation S2.1, *γ* is the swelling rate, *C*_0_ is the concentration of conidia in the experiment, and *σ* is the TAFC secretion rate, the parameter to be estimated. The value of *γ* is known from the model (Table S1) and *C*_0_ from the paper itself. The initial condition is such that TAFC(4) = 0. That is, TAFC is zero at four hours, which comes from Table S1, to be interpreted as conidia starting to swell at four hours. Figure S1 shows the fitting of Eq S2.1 to the experimental data of Hissen, AHT *et al*. 2004 [69]. It should be noted that only *σ* is being fitted, and it should also be noted that the best one can obtain from this experiment is an apparent secretion rate.

**Figure S1:**
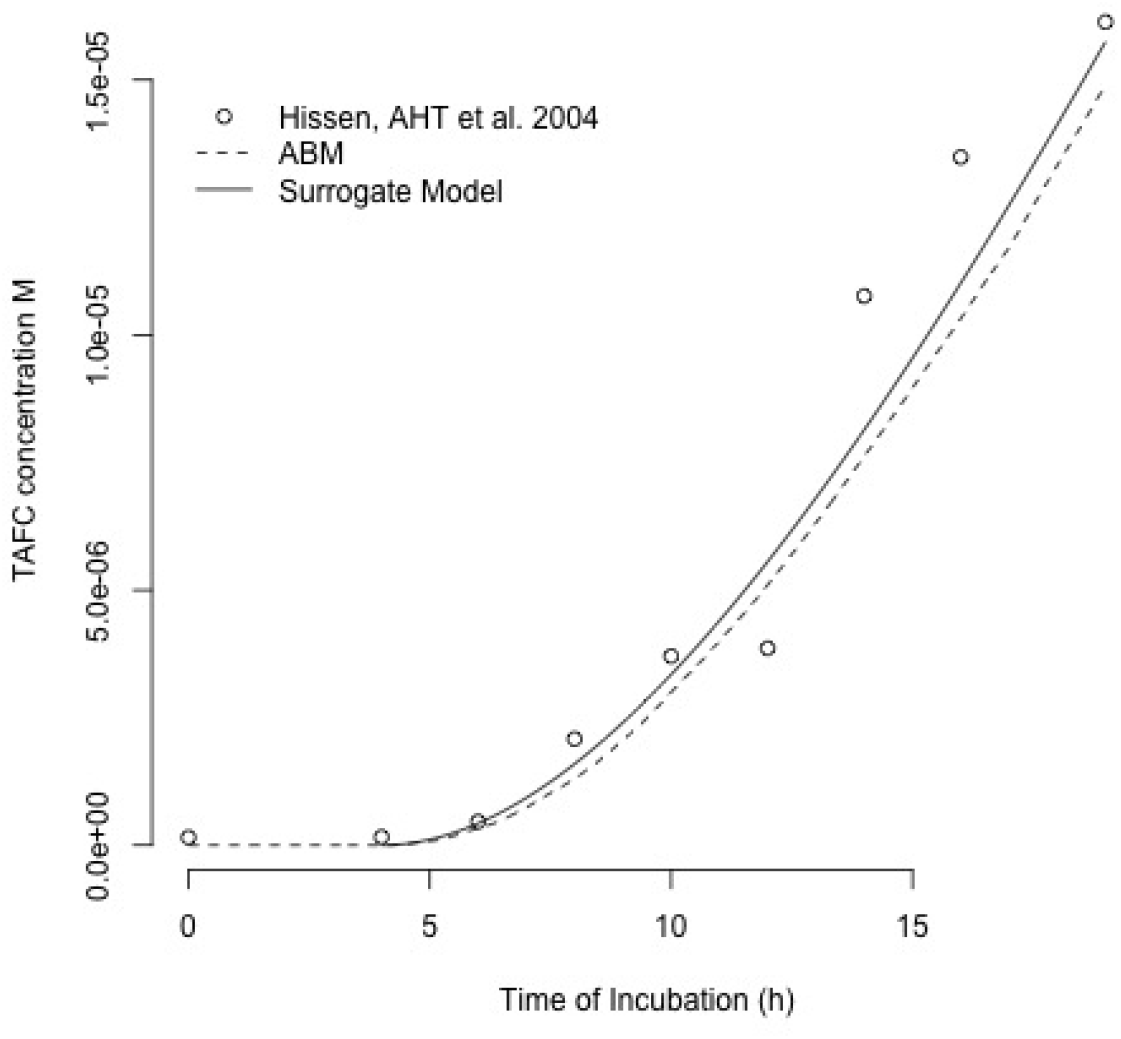
Figure showing the fitting of Eq S2.1. Experimental data from Hissen, AHT *et al*. 2004 [69] (represented as dots in the graph); computed with Eq S2.1 (solid line); computed with the data model (dashed line). Eq S2.1 was computed only from 4h onwards. Before that, we considered no swelling conidia and, therefore, no TAFC production (Table S1). The fact that the full simulator agrees with the surrogate model (Eq S2.1) shows that our procedure was appropriate.

To get the TAFC uptake rate, Raymond-Bouchard, I *et al*. 2012 [70] was used, who reports the TAFC uptake rate as OD600, and Yap, PY *et al*. 2019 [71] reports a curve of OD600 for yeast cells per mL. Supposing that *A. fumigatus* conidia OD600 is similar to that for yeast, one can calculate the TAFC uptake rate per *Aspergillus* cell.

As an approximation, it was assumed that a resting conidium contains one *k*_*d*_ (Section S2.6) of iron, therefore the *Aspergillus* initial amount of iron is *k*_*d*_*_Af _IRON* × *CONIDIA_VOL* (Table S1).

### S2.3 Cytokine and chemokine *k*_*d*_

The *k*_*d*_ of cytokines/chemokines is divided into two distinct sets of data: the *k*_*d*_ of cytokine/chemokine receptors and dose-response curves for these molecules. As an example of a dose-response curve, consider the level of activation of *NF* −*κB* vs. concentration of TNF. It was found that these two approaches are remarkably consistent; that is, activation of a cell by a molecule seems to depend, at least in general, only on the receptor affinity. To model dose-response curves, Equation 2 (Materials & Methods) was used.

### S2.4 Iron dynamics

The IL-6-hepcidin and hepcidin-Tf curves were obtained from the literature. While the macrophage iron import rate was also obtained from the literature, the export rate was assumed to be equal to the import rate. Homeostasis is assumed under normal conditions. Equality of iron import and export are interpreted to mean that the fluxes are equal, not the equations’ constants. Iron import depends on external iron levels while export depends on internal iron levels.

Values for the internal concentration of macrophage iron, transferrin, and saturation are taken from Parmar, JH *et al*. 2019 [66].

### S2.5 Phagocytosis, interaction, and killing rates

A simple law of mass action between leukocytes and conidia is assumed and used to derive Equation, (Eq S2.2). Given time and leukocyte concentration, this equation returns the probability of phagocytosing a conidium. The phagocytosis probability in Table S1 is an extrapolation of Equation S2.2 for a voxel’s local concentration and one time step:

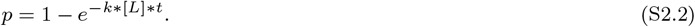

In Eq. S2.2, *p* is the probability of phagocytosis or interaction, [L] is the leukocyte concentration, and *t* is time, taken to be two minutes in the simulator (one time-step). Note that this extrapolation assumes that the limiting factor in the phagocytosis is the direct interaction between leukocyte and conidia and not the spread rate. According to Hoang, AN *et al*. 2013 [106], the leukocyte movement rate can be quite fast.

The hyphae killing rate is determined similarly. In contrast, the internalized conidia killing rate was acquired based on the percentage of internalized conidia killed by macrophages after 12h. Maximum conidia per macrophage is the apparent maximum reported by Gresnigt, MS *et al*. 2018 [87], and for neutrophils, it is a scaling of this number based on the relative size of a neutrophil.

### S2.6 *Aspergillus* iron sensitivity

*Aspergillus* iron sensitivity (*k*_*d*_*_Af _IRON* in Table S1) is a critical parameter in the model. This value measures the concentration of iron needed to turn the sreA gene on/off. This gene, in turn, controls the secretion and uptake of TAFC [107]. As a simplification, this value is also used to control *Aspergillus* growth. Schrettl, M *et al*. 2008 [72] grew WT and sreA KO *Aspergillus* in the presence of TAFCBI (TAFC bound to iron) and then measured the content of iron as *μ* mol per gram of dry weight (DW) in the colonies.

The first thing to notice is that it is safe to assume that both colonies grew to approximately equal size based on Schrettl, M *et al*. 2008, and others. As mentioned, the paper measures iron in *μmol/g* (DW). To convert this to molar, one can first convert DW into wet weight using data from Bakken, LR, 1983 [93]; this paper also gives the fungal density. With that one is able to calculate molarity.

The sreA KO cannot control the influx of TAFC. Therefore, one can assume, for simplicity, that the iron acquisition in these colonies follows a *quasi-linear* equation (Eq S2.3):

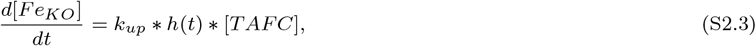

where *k*_*up*_ is the apparent TAFC uptake rate, *h*(*t*) is the equation describing hyphal growth, and [TAFC] is the amount of TAFC in the experiment. This is assumed to be constant, that is, the quantity taken up by hyphae is small compared to the amount supplied. Integrating this equation, one gets:

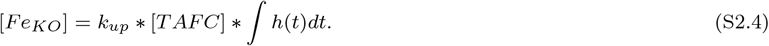

This equation gives the quantity of internal iron at the end of the experiment with the sreA KO *Aspergillus*. For WT colonies, as the iron concentration increases, sreA gets activated, which leads to the downregulation of the TAFC receptor. Therefore, for WT, one has:

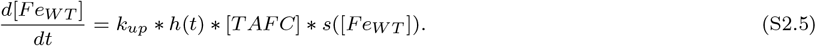

Here, *s*([*Fe*_*WT*_]) is the unknown sreA activation function, but again, one can employ a surrogate model, namely Eq. 2 (Materials & Methods) as a phenomenological model of sreA activation/inactivation. More specifically, this function is used to activate the LIP node that then inactivates the sreA node (see Brandon, M *et al*. 2015 [107]). The function *s*([*Fe*_*WT*_]) should be the complement of that; therefore, Eq S2.5 becomes:

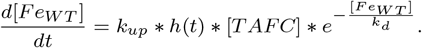

Integrating this equation one obtains:

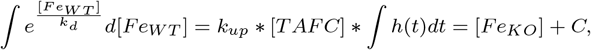

where *C* = *k*_*d*_. Making some algebraic rearrangements results in:

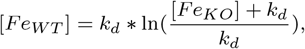

where [*Fe*_*WT*_] is the quantity of iron in the WT experiment, [*Fe*_*KO*_] is the quantity of iron in the sreA KO experiment, and *k*_*d*_ is the value to be determined. The [TAFC] concentration is assumed to be high, compared to the amount consumed and that *h*(*t*) is equal for both sreA KO and WT.

### S2.7 Transferrin Michaelis constant

For convenience, Michaelian kinetics is assumed. The TAFC-Tf kinetics is a complex mechanism described by Hissen, AHT & Moore MM 2005 [67] For convenience, a simplified version is used that does not consider cooperativity or difference in transferrin sites. The reaction rate of lactoferrin is very elusive, not having been studied extensively. Nevertheless, one study injected this protein *in vivo* and saw a 46% decrease in serum iron upon 4h [31]. That evidence enables an educated guess of the reaction rate.

### S2.8 Fungal biology

Fungal dimensions are obtained from the papers cited in Table S1. In some cases, values can be obtained directly from the photomicrograph reported in these papers using the MATLAB App Grabit. Growth rates come from papers that report hyphal length over time, while branching probability was based on the hyphal growth unit length. This gives an estimate of how many branches per septum there are.

A crucial parameter in the model is the swelling rate. Because swelling is quickly succeeded by germination, the germination rate is used as a proxy for the swelling rate. White, LO 1977 [98] report the rate of germination *in vivo*, and Gago, S *et al*. 2018 report a very consistent value *in vitro* in the presence of bronchoepithelial cells. From these papers, one can make a robust estimate of this value.

### S2.9 Number of cells and lung size

Calculations were made by considering a pair of inflated mouse lungs, which has a volume of 1mL [108]. We consider that these pairs of lungs have around 3 million alveoli [109, 110], 230,000 resident macrophages [104], and ten million type-II epithelial cells [105]. The maximum number of mono- and polymorphonuclear cells come from a paper reporting counts of these leukocytes per hundred alveoli in mice infected with *Aspergillus* [103]. The global recruitment rate was adjusted to fit the curve of neutrophils in Bonett, CR *et al*. 2006 [104]. Note that the numbers of macrophages and neutrophils reflect those of bronchoalveolar lavage fluid.

### S2.10 Turnover rate

To calculate the turnover rate between the lung and the rest of the body, the following system of differential equations was used:

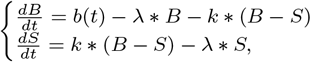

where *B* is the concentration of the molecule in BAL, *S* is the concentration in serum, and *λ* is the decay rate. The function *b*(*t*) is the secretion rate, and *k* is the exchange rate, the value to be estimated. The ratio of interest is *B/S* in the equilibrium. Notice that upon algebraic rearrangements, one finds that this ratio is (*k* + *λ*)*/k*. But *λ* is known from the literature (Table S1), and one can estimate *k* from an empirical *B/S* ratio found in Gonçalves, SM *et al*. 2017 [102].

### S2.11 Leukocyte movement

This movement rate can be obtained from Khandoga, AG *et al*. 2009 [79]. The value, 1.44 *μm/min*, is conservative compared to other sources. Pollmacher J, & Figge MT, 2014 [111] uses a movement rate of 2-6 *μm/min*, for instance. Nevertheless, the rate used here must be considered a phenomenological movement rate. In the real lung, leukocytes may not move in a straight line but along the alveolar curved surface. That is the case in the Pollmacher J, & Figge MT, 2014 [111] model.

### S2.12 Antibodies

Antibody parameters were not specific for TNF. The concentration was obtained based on a measurement of *IgG* found in mice after an immunization assay. The value of *k*_*cat*_ and *Km* resulted from the imposition of Michaelian kinetics, using data for a generic protein antigen, in this case, the lysosome. Half-life comes from Vieira, P, and Rajewsky K 1988 [63].

### S2.13 Other parameters

The time that cells need to change status (T_CHANGE and T_REST) were based on *in-vitro* reports [99]. The half-life of molecules is an average. Likewise, the diffusion rate is an average of the values of the different molecules in living tissue.

For macrophages/monocytes, the 24h value reported for monocytes [91] is used. Other literature sources nevertheless report a longer half-life for macrophages. To calculate macrophage volume, they can be approximated by a sphere, and one can then use the dimensions reported by Krombach, F *et al*. 1997 [100].

## S3 Interaction Rules

**Table S2:**
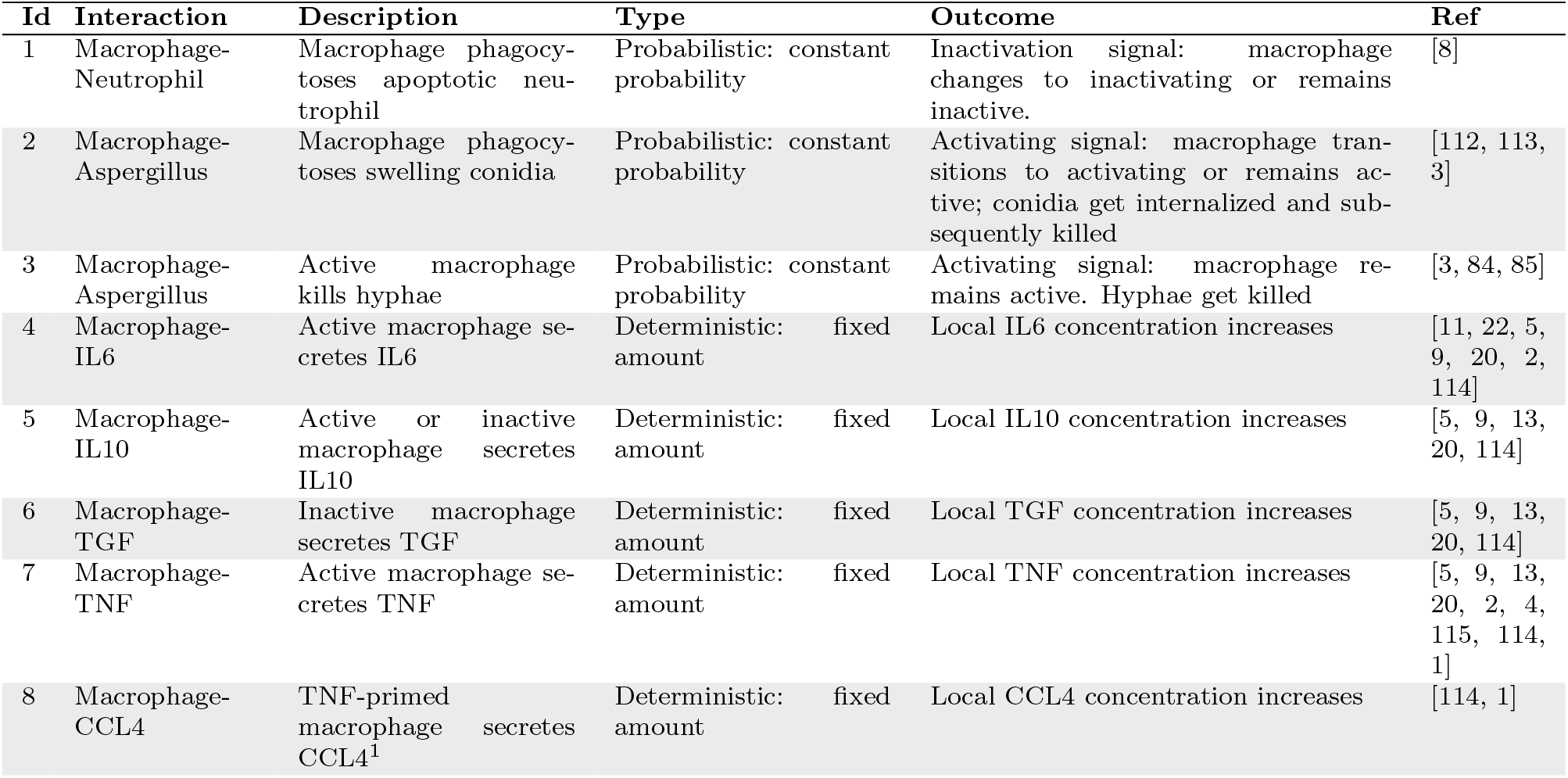
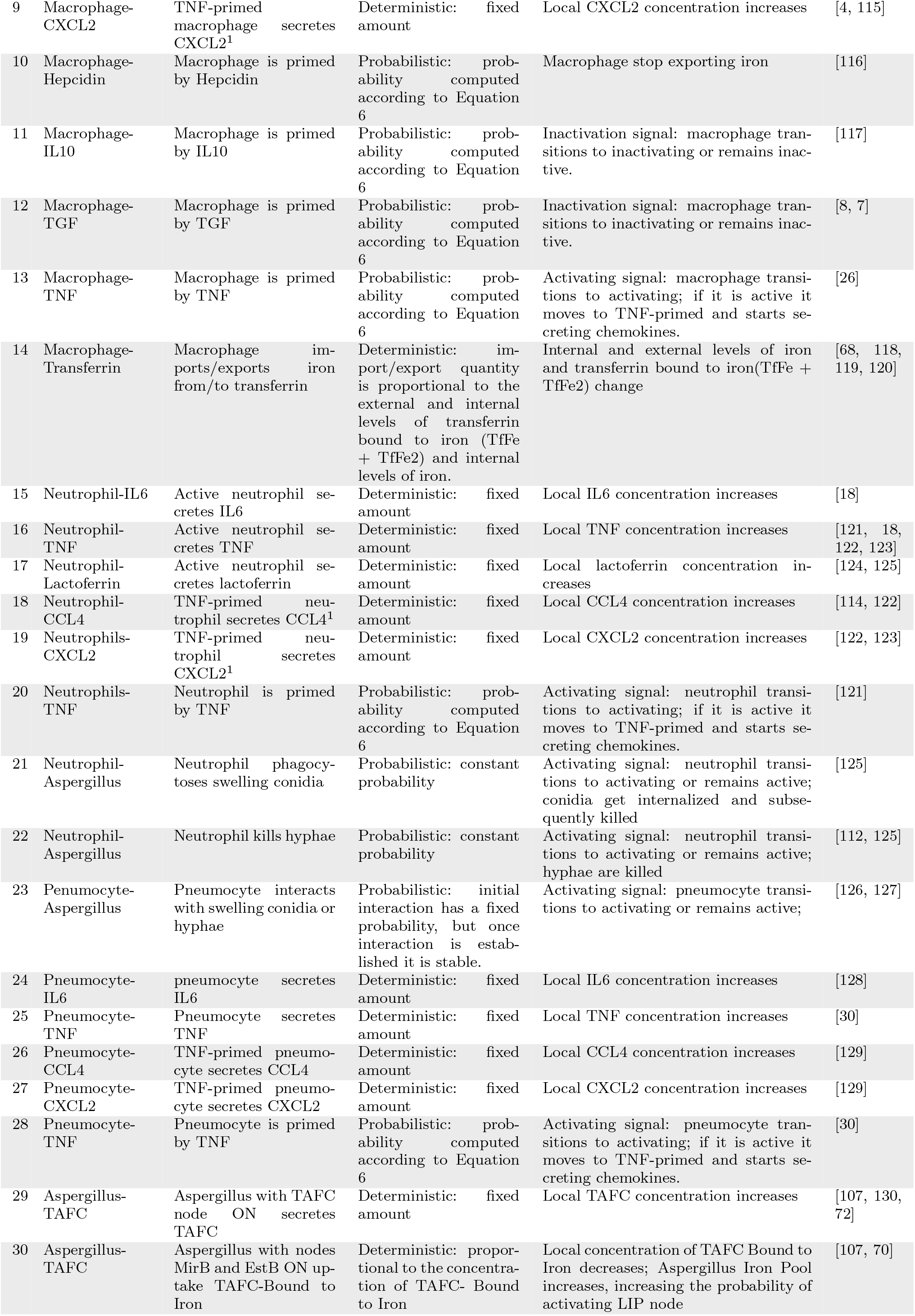
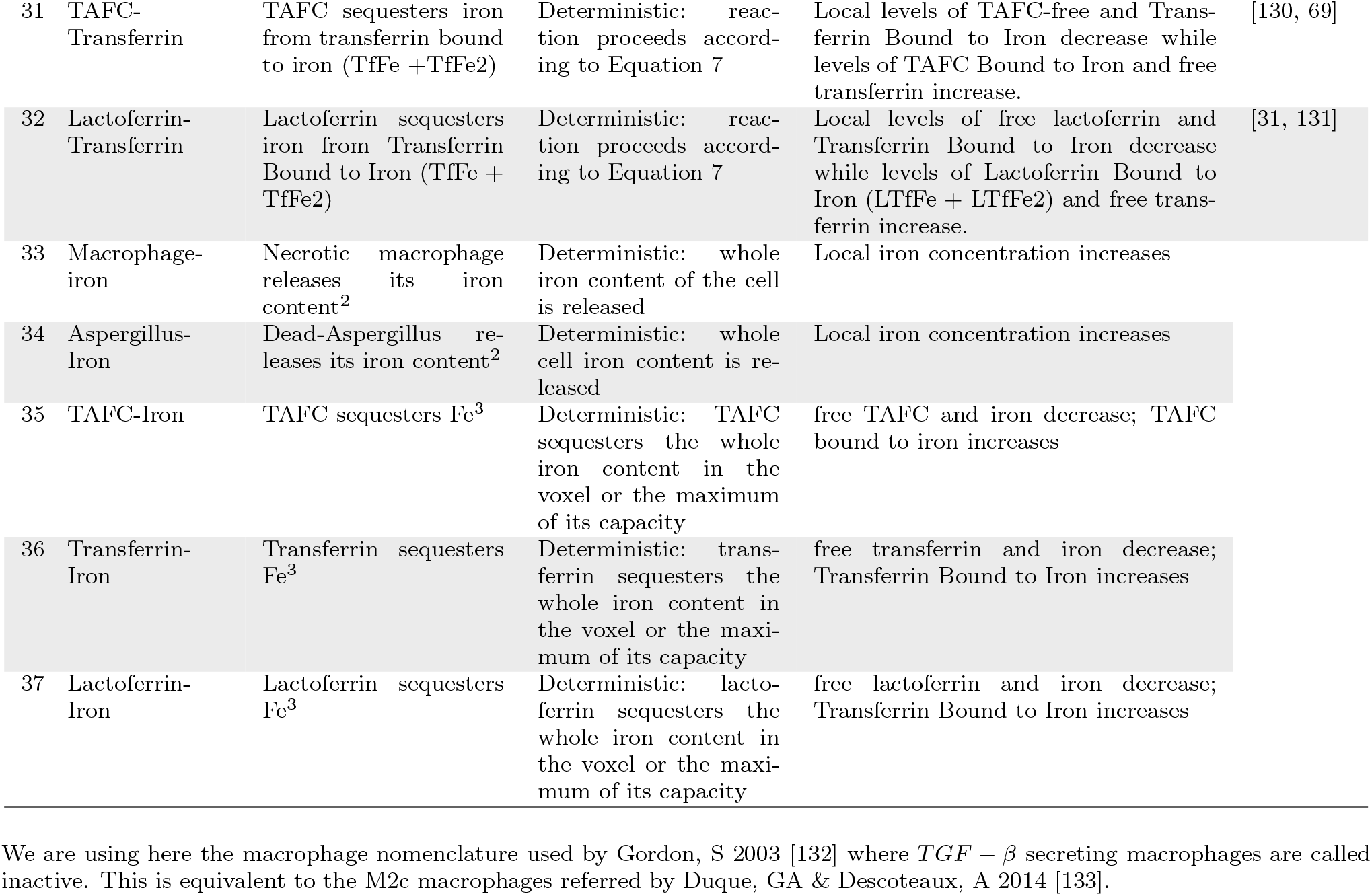
Table with interaction rules. Column one shows the two agents or molecules involved in the interaction. Column three briefly describes the interaction. Column four describes if the interaction is deterministic or probabilistic and if there is an equation involved. Column five describes the outputs of the interaction, that is, how the agent’s states are modified upon the interactions and how the concentration of the molecules changes upon the interaction. Notice that an agent can interact with another agent or molecule in more than one way. Therefore, there is some duplication in column two. However, this duplication is eliminated upon the description of the interaction.

## S4 Drivers of Fungal Burden

**Figure S2:**
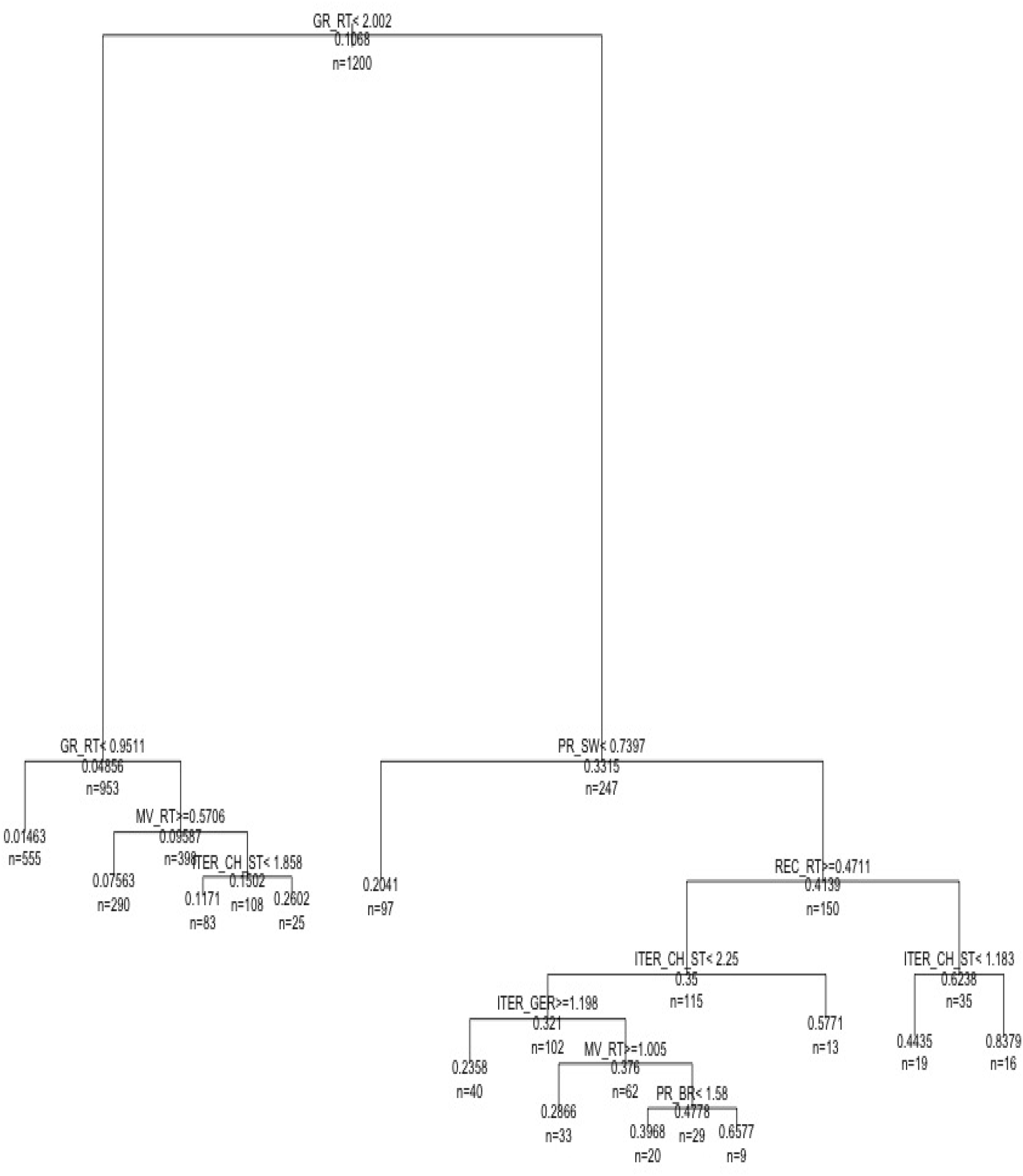
Classification trees of fungal burden. We generated 1200 samples with LHS and ran simulations with immunocompetent hosts, and classified the simulations by the maximum number of hyphal cells measured over 24 hours of simulation. For convenience, all the parameters were normalized by their default values. That is, a value of 2 means that the tree made a partition when that parameter was twice its default value (Table S1). We also normalized the fungal burden by the initial dose inocula (1920 conidia). Notice, however, that the measure of fungal burden is the maximum number of hyphal cells encountered at any time over a 24 hour period. This counts only the conidia that germinate and the hyphae that multiply, excluding ungerminated conidia.

**Figure S3:**
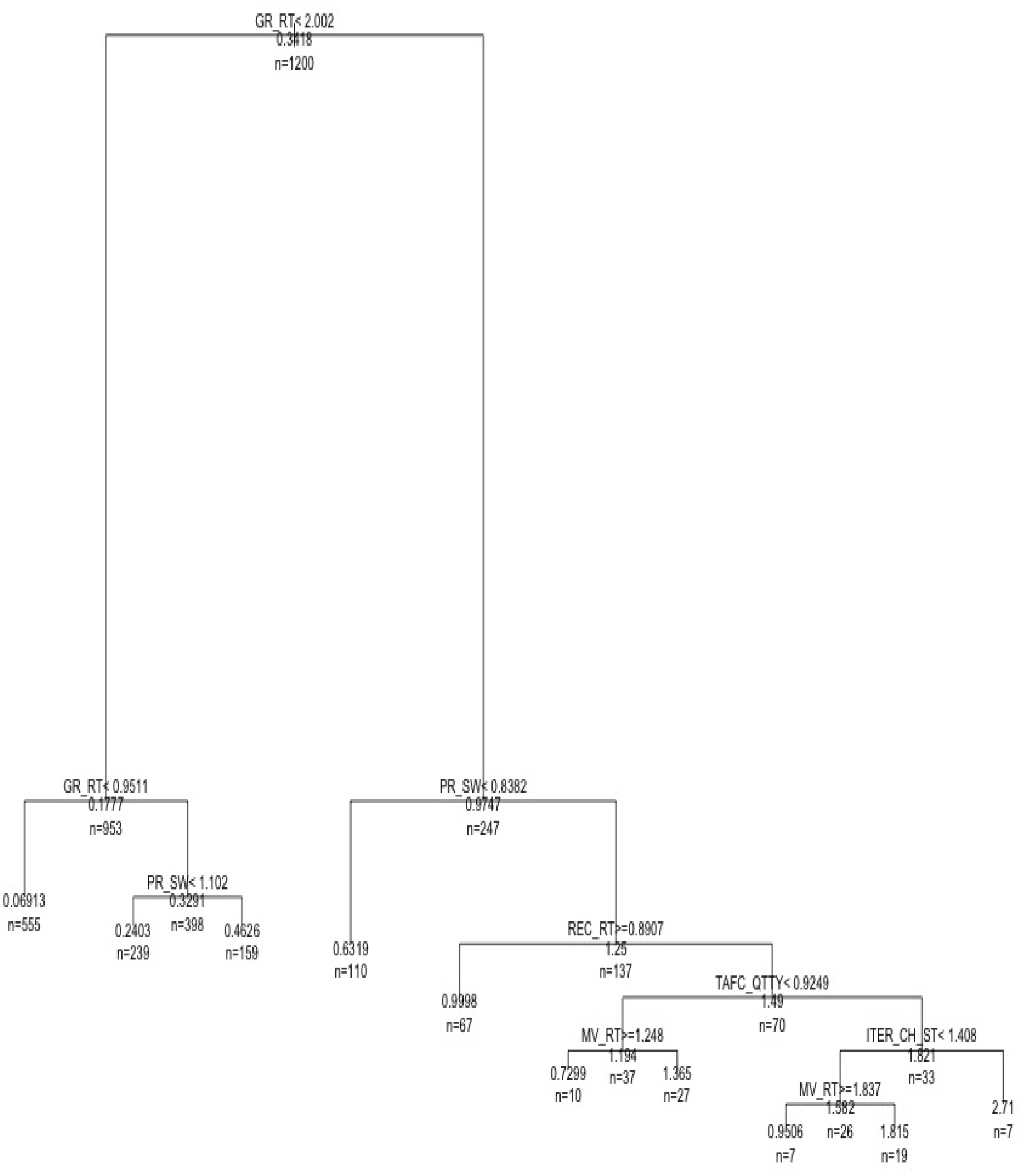
Classification trees of fungal burden. We generated 1200 samples with LHS and ran simulations with neutropenic hosts, and classified the simulations by the maximum number of hyphal cells measured over 24 hours of simulation. For convenience, all the parameters were normalized by their default values. That is, a value of 2 means that the tree made a partition when that parameter was twice its default value (Table S1). We also normalized the fungal burden by the initial dose inocula (1920 conidia). Notice, however, that the measure of fungal burden is the maximum number of hyphal cells encountered at any time over a 24 hour period. This counts only the conidia that germinate and the hyphae that multiply, excluding ungerminated conidia.

**Figure S4:**
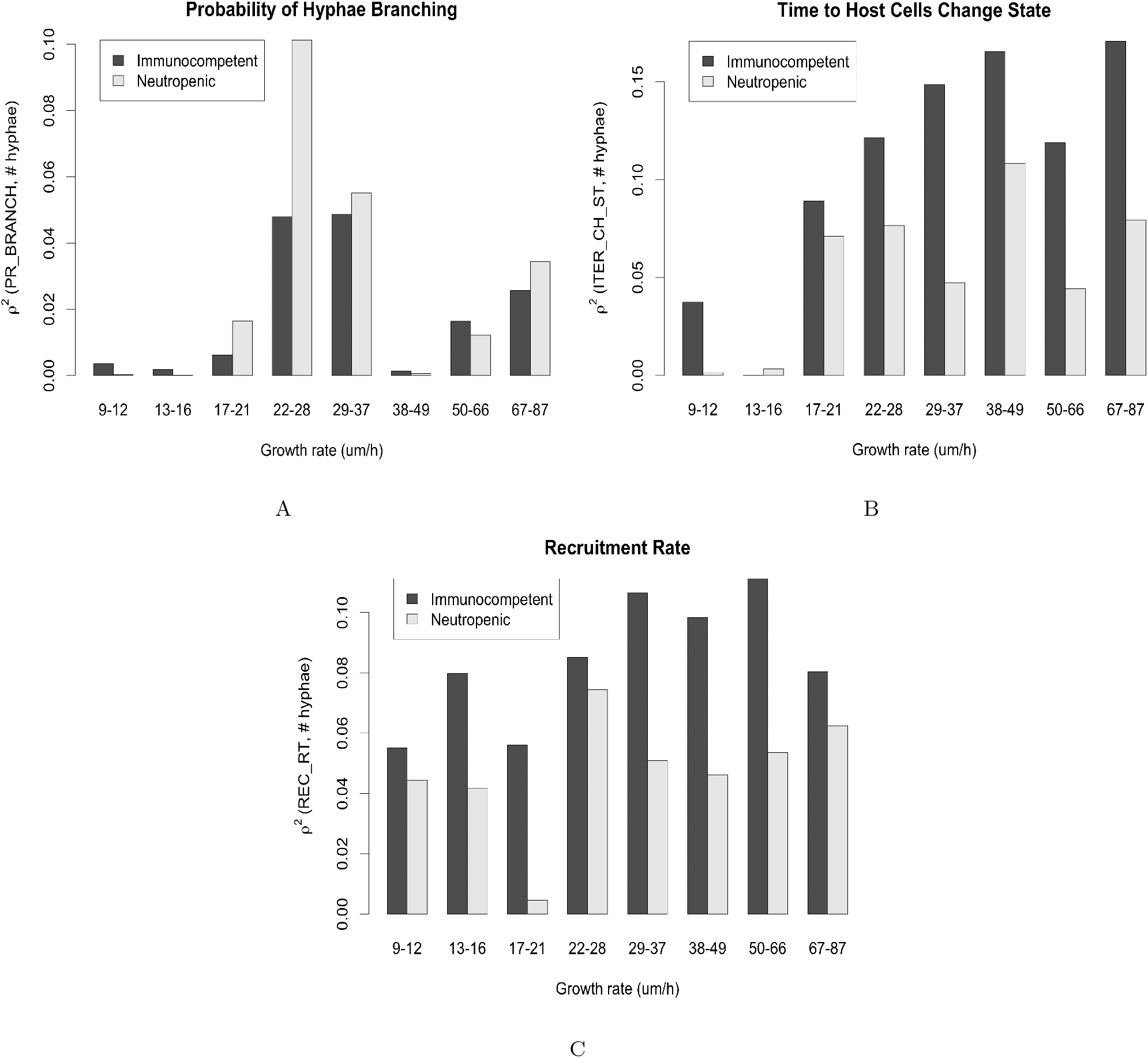
Variation of the square of the correlation *r*^2^ between the critical model parameters and fungal burden, as the growth rate varies along the x-axis. We started with a dataset of 1200 samples generated with LHS and then partitioned this dataset by growth rate. The x-axis in the figure shows the range of the growth rate in each partition. We then calculate the correlation between parameters and fungal burden in each partition. Fungal burden was the maximum number of hyphal cells measured over the course of 24 hours of simulation. A: variation of *r*^2^ between the probability of branching and fungal burden. B: variation of *r*^2^ between the iterations for host cells to change state and fungal burden. C: variation of *r*^2^ between the leukocyte recruitment rate and fungal burden.

## Notes

### Competing Interest Statement

The authors have declared no competing interest.

