## Supplementary File for "Multiscale mechanistic modelling of the host defense in invasive aspergillosis reveals leukocyte activation and iron acquisition as drivers of infection outcome"

Henrique AL Ribeiro, et al.

October 29, 2021

#### S1 The Parameters

Table S1: Table with the model parameters. Parameters 1-14 were obtained in a unified manner as described in the manuscript. As mentioned in the manuscript, "macrophage" should be interpreted as "macrophage/monocyte." Probabilities of phagocytosis, killing, and interaction refer to the likelihood of an event succeeding in one iteration if the appropriate conditions apply. The maximum number of cells and the average number of epithelial cells is computed over the whole simulated space.

| Id | Parameter | Description | Value | Reference |
|---|---|---|---|---|
| 1 | $MA\_IL6\_QTTY$ | Macrophage/monocyte IL6 secretion rate. | $1.46 \times 10^{-20}$ $mol * cell^{-1} * h^{-1}$ | [1, 2, 3, 4, 5, 6, 7, 8, 9, 10, 11, 12, 13, 14, 15, 16, 17, 18, 19, 20, 21, 22, 23, 24, 25, 26, 27, 28, 29, 30] |
| 2 | $MA\_CCL4\_QTTY$ | Macrophage/monocyte CCL4 secretion rate | $1.79 \times 10^{-20}$ $mol * cell^{-1} * h^{-1}$ | |
| 3 | $MA\_CXCL2\_QTTY$ | Macrophage/monocyte CXCL2 secretion rate. | $1.11 \times 10^{-19}$ $mol * cell^{-1} * h^{-1}$ | |
| 4 | $MA\_IL10\_QTTY$ | Macrophage/monocyte IL10 secretion rate. | $6.97 \times 10^{-22}$ $mol * cell^{-1} * h^{-1}$ | |
| 5 | $MA\_TNF\_QTTY$ | Macrophage/monocyte TNF secretion rate. | $3.22 \times 10^{-20}$ $mol * cell^{-1} * h^{-1}$ | |
| 6 | $MA\_TGF\_QTTY$ | Macrophage/monocyte $TGF - \beta$ secretion rate. | $1.01 \times 10^{-21}$ $mol * cell^{-1} * h^{-1}$ | |
| 7 | $N\_IL6\_QTTY$ | Neutrophil IL6 secretion rate. | $8.59 \times 10^{-23}$ $mol * cell^{-1} * h^{-1}$ | |
| 8 | $N\_CXCL2\_QTTY$ | Neutrophil CXCL2 secretion rate. | $6.50 \times 10^{-22}$ $mol * cell^{-1} * h^{-1}$ | |
| 9 | $N\_TNF\_QTTY$ | Neutrophil TNF secretion rate. | $1.89 \times 10^{-22}$ $mol * cell^{-1} * h^{-1}$ | |
| 10 | $E\_IL6\_QTTY$ | Epithelial cell IL6 secretion rate. | $1.46 \times 10^{-20}$ $mol * cell^{-1} * h^{-1}$ | |
| 11 | $E\_CCL4\_QTTY$ | Epithelial cell CCL4 secretion rate | $1.79 \times 10^{-20}$ $mol * cell^{-1} * h^{-1}$ | |
| 12 | $E\_CXCL2\_QTTY$ | Epithelial cell CXCL2 secretion rate. | $1.11 \times 10^{-19}$ $mol * cell^{-1} * h^{-1}$ | |
| 13 | $E\_TNF\_QTTY$ | Epithelial cell TNF secretion rate. | $3.22 \times 10^{-20}$ $mol * cell^{-1} * h^{-1}$ | |
| 14 | $LAC\_QTTY$ | Lactoferrin secretion rate (Neutrophils) | $4.37 \times 10^{-17}$ $mol * cell^{-1} * h^{-1}$ | [31] |
| 15 | $k_d\_IL6$ | IL-6 $k_d$ | 330 pM | [32, 33, 34] |
| 16 | $k_d\_CCL4$ | CCL4 $k_d$ | 180 pM | [35] |
| 17 | $k_d\_CXCL2$ | CXCL2 $k_d$ | 91.667 pM | [36, 37] |
| 18 | $k_d\_IL10$ | IL-10 $k_d$ | 140 pM | [38, 39, 40, 41] |
| 19 | $k_d\_TNF$ | TNF $k_d$ | 326 pM | [42, 43, 44, 45, 46, 47, 48, 49] |
| 20 | $k_d\_TGF - \beta$ | $TGF - \beta$ $k_d$ | 26.5 pM | [50, 51, 52] |

| | | | | |
|---|---|---|---|---|
| 21 | $k_d\_HEP$ | Hepcidin $k_d$ | 855 nM | [53] |
| 22 | $D$ | Diffusion rate | 850 $\mu m^2/min$ | [54, 55] |
| 23 | $\lambda$ | Cytokine/chemokine half-life | 1h | [56, 57, 58, 59, 60, 61, 62] |
| 24 | $\lambda_{ab}$ | Antibody half-life (Anti-TNF) | 5 days | [63] |
| 25 | $HEP\_INT$ | Hepcidin intercept (IL6-hepcidin model) | -0.3141 | [64] |
| 26 | $HEP\_SLOPE$ | Hepcidin slope (IL6-hepcidin model) | 0.78 | [64] |
| 27 | $Tf\_INT$ | Transferrin intercept (Tf-hepcidin model) | $-1.194 \times 10^{-5}$ | [65] |
| 28 | $Tf\_SLOPE$ | Transferrin slope (Tf-hepcidin model) | $-5.523 \times 10^{-6}$ | [65] |
| 29 | $Def\_TF\_CON.$ | Default Tf concentration | 32.25 $\mu M$ | [66, 64, 65] |
| 30 | $APO\_Tf\_REL\_CON$ | Apo-Tf relative concentration | 40% | [66, 67] |
| 31 | $TfFe\_REL\_CON$ | Monoferric Transferrin relative concentration | 16.57% | |
| 32 | $TfFe2\_REL\_CON$ | Diferric Transferrin relative concentration | 43.43% | |
| 33 | $MA\_IRON\_EXP$ | Macrophage iron export rate | $1367.30 M^{-1} * h^{-1}$ | [68] |
| 34 | $MA\_IRON\_IMP$ | Macrophage iron uptake rate | $5.33 \times 10^{-12} L * cell^{-1} * h^{-1}$ | [68] |
| 35 | $MA\_INT\_IRON$ | Macrophage initial internal iron quantity | $1.0086 \times 10^{-14} mol$ | [66] |
| 36 | $TAFC\_QTTY$ | TAFC secretion rate. | $1.0 \times 10^{-15}$ $mol * cell^{-1} * h^{-1}$ | [69] |
| 37 | $TAFCBI\_UPTAKE$ | TAFCBI (Bound to Iron) uptake rate | $1 \times 10^{-12} L * cell^{-1} * h^{-1}$ | [70, 71] |
| 38 | $k_d\_Af\_IRON$ | *A. fumigatus* iron sensitivity | 79.05 $\mu M$ | [72] |
| 39 | $K_{cat}/K_M\_TAFC$* | $K_{cat}/K_M$ TAFC-Tf | 397.77 $M^{-1}s^{-1}$ | [69, 67] |
| 40 | $K_{cat}/K_M\_LAC$* | $K_{cat}/K_M$ Lactoferrin-Tf. | 399.2 $M^{-1}s^{-1}$ | [31] |
| 41 | $K_{cat}\_AB$ | $K_{cat}$ Antibody-Antigen | 120 $s^{-1}$ | [73] |
| 42 | $K_M\_AB$ | $K_M$ Antibody-Antigen | 0.697 $\mu M$ | [73] |
| 43 | $AB\_CON$ | Antibody systemic concentration upon injection | 0.2 nM | [74] |
| 44 | $r$ | *A. fumigatus* growth rate | 41.3 $\mu m/h$ | [75, 76, 77, 78] |
| 45 | $MV\_RT$ | Leukocyte movement rate | 1.44 $\mu m/min$ | [79] |
| 46 | $N\_H\_KILL$ | Neutrophil-hyphae killing probability | 22.71%. | [80, 81, 82, 83] |
| 47 | $MA\_H\_KILL$ | Macrophage-hyphae killing probability. | 9.85% | [84, 85] |
| 48 | $MA\_PHAG$ | Macrophage phagocytosis probability | 90.55% | [86, 87] |
| 49 | $N\_PHAG$ | Neutrophil phagocytosis probability | 14.73% | [83] |
| 50 | $MA\_MAX\_CONIDIA$ | Max ingested conidia by macrophage | 18 | [87] |
| 51 | $N\_MAX\_CONIDIA$ | Max ingested conidia by neutrophils | 3 | [87, 88] |
| 52 | $E\_INT$ | Resting epithelial cell-*Aspergillus* interaction probability | 4.49% | [89] |
| 53 | $PHAG\_KILL$ | Probability of killing internalized conidia (leukocytes) | 1.28% | [90] |

| | | | | |
|---|---|---|---|---|
| 54 | $MA\_HALF\_LIFE$ | Macrophage/monocyte half-life | 24h | [91] |
| 55 | $N\_HALF\_LIFE$ | Neutrophil half-life | 6h | [92] |
| 56 | $SEPTAE\_r$ | Septae radius | 2.65 $\mu m$ | [93, 94, 95] |
| 57 | $SEPTAE\_L$ | Septae length | 40 $\mu m$ | [95, 96] |
| 58 | $T\_SWELL$ | Time to start swelling | 4h | [97] |
| 59 | $PR\_SW$ | Swelling probability | 0.39% | [77, 98] |
| 60 | $ITER\_GER$ | Time until germination (after swelling) | 2h | [97] |
| 61 | $ITER\_CH\_ST$ | Iterations to cell state change | 60 | [99] |
| 62 | $INTER\_REST$ | Iterations until active cells return to resting | 180 | [99] |
| 63 | $H\_VOL$ | Hyphae volume | 1.06 pL | [93, 94, 95, 96] |
| 64 | $MA\_VOL$ | Macrophage volume | 4.85 pL | [100] |
| 65 | $CONIDIA\_VOL$ | Conidia volume | 0.0484 pL | [94] |
| 66 | $PR\_BR$ | Probability of branching (*A. fumigatus*) | 25% | [101, 75] |
| 67 | $TURNOVER\_RT$ | Molecule exchange rate between lung and whole body serum. | 0.1823 $h^{-1}$ | [102] |
| 68 | $MAX\_N$ | Maximum number of neutrophils | 522 | [103] |
| 69 | $MIN\_N$ | Minimum number of neutrophils | 0 | [104] |
| 70 | $MAX\_MA$ | Maximum number of macrophages | 209 | [103] |
| 71 | $MIN\_MA$ | Minimum number of macrophages | 15 | [104] |
| 72 | $AVG\_E$ | Average number of epithelial cells | 640 | [105] |
| 73 | $REC\_RT$ | Global recruitment rate | 2 | [104] |
| 74 | $Af\_INIT\_IRON$ | *A. fumigatus* initial iron. | $3.83 \times 10^{-18} mol$ | [94, 72] |

$$\frac{d^2[TAFC]}{dt^2} + \gamma * \frac{d[TAFC]}{dt} = \gamma * \sigma * C_0. \quad \text{(S2.1)}$$

In Equation S2.1, $\gamma$ is the swelling rate, $C_0$ is the concentration of conidia in the experiment, and $\sigma$ is the TAFC secretion rate, the parameter to be estimated. The value of $\gamma$ is known from the model (Table S1) and $C_0$ from the paper itself. The initial condition is such that TAFC(4) = 0. That is, TAFC is zero at four hours, which comes from Table S1, to be interpreted as conidia starting to swell at four hours. Figure S1 shows the fitting of Eq S2.1 to the experimental data of Hissen, AHT *et al.* 2004 [69]. It should be noted that only $\sigma$ is being fitted, and it should also be noted that the best one can obtain from this experiment is an apparent secretion rate.

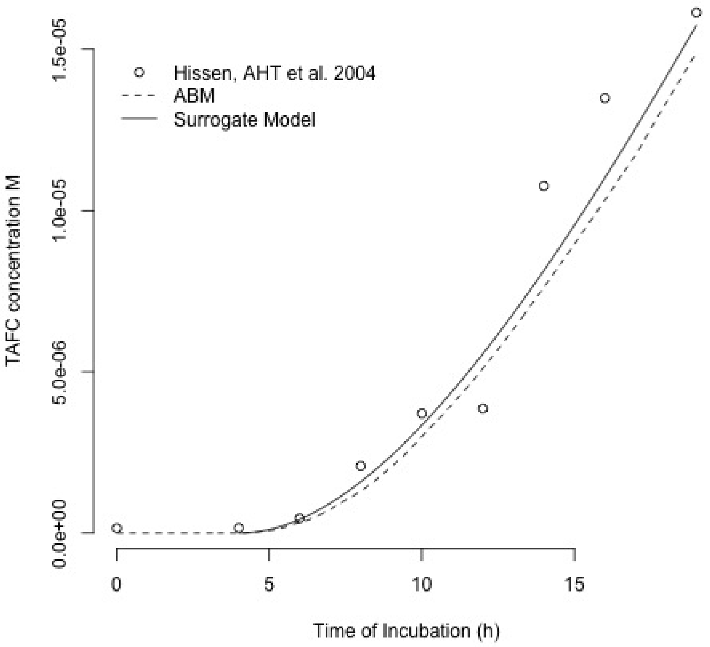


Figure S1: Figure showing the fitting of Eq S2.1. Experimental data from Hissen, AHT *et al.* 2004 [69] (represented as dots in the graph); computed with Eq S2.1 (solid line); computed with the data model (dashed line). Eq S2.1 was computed only from 4h onwards. Before that, we considered no swelling conidia and, therefore, no TAFC production (Table S1). The fact that the full simulator agrees with the surrogate model (Eq S2.1) shows that our procedure was appropriate.

The sreA KO cannot control the influx of TAFC. Therefore, one can assume, for simplicity, that the iron acquisition in these colonies follows a *quasi-linear* equation (Eq S2.3):

$$\frac{d[Fe_{KO}]}{dt} = k_{up} * h(t) * [TAFC], \tag{S2.3}$$

where $k_{up}$ is the apparent TAFC uptake rate, $h(t)$ is the equation describing hyphal growth, and [TAFC] is the amount of TAFC in the experiment. This is assumed to be constant, that is, the quantity taken up by hyphae is small compared to the amount supplied. Integrating this equation, one gets:

$$[Fe_{KO}] = k_{up} * [TAFC] * \int h(t)dt. \tag{S2.4}$$

This equation gives the quantity of internal iron at the end of the experiment with the sreA KO *Aspergillus*. For WT colonies, as the iron concentration increases, sreA gets activated, which leads to the downregulation of the TAFC receptor. Therefore, for WT, one has:

$$\frac{d[Fe_{WT}]}{dt} = k_{up} * h(t) * [TAFC] * s([Fe_{WT}]). \tag{S2.5}$$

Here, $s([Fe_{WT}])$ is the unknown sreA activation function, but again, one can employ a surrogate model, namely Eq. 2 (Materials & Methods) as a phenomenological model of sreA activation/inactivation. More specifically, this function is used to activate the LIP node that then inactivates the sreA node (see Brandon, M *et al.* 2015 [107]). The function $s([Fe_{WT}])$ should be the complement of that; therefore, Eq S2.5 becomes:

$$\frac{d[Fe_{WT}]}{dt} = k_{up} * h(t) * [TAFC] * e^{-\frac{[Fe_{WT}]}{k_d}}.$$

Integrating this equation one obtains:

$$\int e^{\frac{[Fe_{WT}]}{k_d}} d[Fe_{WT}] = k_{up} * [TAFC] * \int h(t)dt = [Fe_{KO}] + C,$$

where $C = k_d$. Making some algebraic rearrangements results in:

$$[Fe_{WT}] = k_d * \ln(\frac{[Fe_{KO}] + k_d}{k_d}),$$

#### S2.10 Turnover rate

To calculate the turnover rate between the lung and the rest of the body, the following system of differential equations was used:

$$\begin{cases} \frac{dB}{dt} = b(t) - \lambda * B - k * (B - S) \\ \frac{dS}{dt} = k * (B - S) - \lambda * S, \end{cases}$$

### S3 Interaction Rules

Table S2: Table with interaction rules. Column one shows the two agents or molecules involved in the interaction. Column three briefly describes the interaction. Column four describes if the interaction is deterministic or probabilistic and if there is an equation involved. Column five describes the outputs of the interaction, that is, how the agent's states are modified upon the interactions and how the concentration of the molecules changes upon the interaction. Notice that an agent can interact with another agent or molecule in more than one way. Therefore, there is some duplication in column two. However, this duplication is eliminated upon the description of the interaction.

| Id | Interaction | Description | Type | Outcome | Ref |
|---|---|---|---|---|---|
| 1 | Macrophage-Neutrophil | Macrophage phagocytoses apoptotic neutrophil | Probabilistic: constant probability | Inactivation signal: macrophage changes to inactivating or remains inactive. | [8] |
| 2 | Macrophage-Aspergillus | Macrophage phagocytoses swelling conidia | Probabilistic: constant probability | Activating signal: macrophage transitions to activating or remains active; conidia get internalized and subsequently killed | [112, 113, 3] |
| 3 | Macrophage-Aspergillus | Active macrophage kills hyphae | Probabilistic: constant probability | Activating signal: macrophage remains active. Hyphae get killed | [3, 84, 85] |
| 4 | Macrophage-IL6 | Active macrophage secretes IL6 | Deterministic: fixed amount | Local IL6 concentration increases | [11, 22, 5, 9, 20, 2, 114] |
| 5 | Macrophage-IL10 | Active or inactive macrophage secretes IL10 | Deterministic: fixed amount | Local IL10 concentration increases | [5, 9, 13, 20, 114] |
| 6 | Macrophage-TGF | Inactive macrophage secretes TGF | Deterministic: fixed amount | Local TGF concentration increases | [5, 9, 13, 20, 114] |
| 7 | Macrophage-TNF | Active macrophage secretes TNF | Deterministic: fixed amount | Local TNF concentration increases | [5, 9, 13, 20, 2, 4, 115, 114, 1] |
| 8 | Macrophage-CCL4 | TNF-primed macrophage secretes CCL4[1] | Deterministic: fixed amount | Local CCL4 concentration increases | [114, 1] |

| | | | | | |
|---|---|---|---|---|---|
| 9 | Macrophage-CXCL2 | TNF-primed macrophage secretes CXCL2[1] | Deterministic: fixed amount | Local CXCL2 concentration increases | [4, 115] |
| 10 | Macrophage-Hepcidin | Macrophage is primed by Hepcidin | Probabilistic: probability computed according to Equation 6 | Macrophage stop exporting iron | [116] |
| 11 | Macrophage-IL10 | Macrophage is primed by IL10 | Probabilistic: probability computed according to Equation 6 | Inactivation signal: macrophage transitions to inactivating or remains inactive. | [117] |
| 12 | Macrophage-TGF | Macrophage is primed by TGF | Probabilistic: probability computed according to Equation 6 | Inactivation signal: macrophage transitions to inactivating or remains inactive. | [8, 7] |
| 13 | Macrophage-TNF | Macrophage is primed by TNF | Probabilistic: probability computed according to Equation 6 | Activating signal: macrophage transitions to activating; if it is active it moves to TNF-primed and starts secreting chemokines. | [26] |
| 14 | Macrophage-Transferrin | Macrophage imports/exports iron from/to transferrin | Deterministic: import/export quantity is proportional to the external and internal levels of transferrin bound to iron (TfFe + TfFe2) and internal levels of iron. | Internal and external levels of iron and transferrin bound to iron(TfFe + TfFe2) change | [68, 118, 119, 120] |
| 15 | Neutrophil-IL6 | Active neutrophil secretes IL6 | Deterministic: fixed amount | Local IL6 concentration increases | [18] |
| 16 | Neutrophil-TNF | Active neutrophil secretes TNF | Deterministic: fixed amount | Local TNF concentration increases | [121, 18, 122, 123] |
| 17 | Neutrophil-Lactoferrin | Active neutrophil secretes lactoferrin | Deterministic: fixed amount | Local lactoferrin concentration increases | [124, 125] |
| 18 | Neutrophil-CCL4 | TNF-primed neutrophil secretes CCL4[1] | Deterministic: fixed amount | Local CCL4 concentration increases | [114, 122] |
| 19 | Neutrophils-CXCL2 | TNF-primed neutrophil secretes CXCL2[1] | Deterministic: fixed amount | Local CXCL2 concentration increases | [122, 123] |
| 20 | Neutrophils-TNF | Neutrophil is primed by TNF | Probabilistic: probability computed according to Equation 6 | Activating signal: neutrophil transitions to activating; if it is active it moves to TNF-primed and starts secreting chemokines. | [121] |
| 21 | Neutrophil-Aspergillus | Neutrophil phagocytoses swelling conidia | Probabilistic: constant probability | Activating signal: neutrophil transitions to activating or remains active; conidia get internalized and subsequently killed | [125] |
| 22 | Neutrophil-Aspergillus | Neutrophil kills hyphae | Probabilistic: constant probability | Activating signal: neutrophil transitions to activating or remains active; hyphae are killed | [112, 125] |
| 23 | Penumocyte-Aspergillus | Pneumocyte interacts with swelling conidia or hyphae | Probabilistic: initial interaction has a fixed probability, but once interaction is established it is stable. | Activating signal: pneumocyte transitions to activating or remains active; | [126, 127] |
| 24 | Pneumocyte-IL6 | pneumocyte secretes IL6 | Deterministic: fixed amount | Local IL6 concentration increases | [128] |
| 25 | Pneumocyte-TNF | Pneumocyte secretes TNF | Deterministic: fixed amount | Local TNF concentration increases | [30] |
| 26 | Pneumocyte-CCL4 | TNF-primed pneumocyte secretes CCL4 | Deterministic: fixed amount | Local CCL4 concentration increases | [129] |
| 27 | Pneumocyte-CXCL2 | TNF-primed pneumocyte secretes CXCL2 | Deterministic: fixed amount | Local CXCL2 concentration increases | [129] |
| 28 | Pneumocyte-TNF | Pneumocyte is primed by TNF | Probabilistic: probability computed according to Equation 6 | Activating signal: pneumocyte transitions to activating; if it is active it moves to TNF-primed and starts secreting chemokines. | [30] |
| 29 | Aspergillus-TAFC | Aspergillus with TAFC node ON secretes TAFC | Deterministic: fixed amount | Local TAFC concentration increases | [107, 130, 72] |
| 30 | Aspergillus-TAFC | Aspergillus with nodes MirB and EstB ON uptake TAFC-Bound to Iron | Deterministic: proportional to the concentration of TAFC- Bound to Iron | Local concentration of TAFC Bound to Iron decreases; Aspergillus Iron Pool increases, increasing the probability of activating LIP node | [107, 70] |

| | | | | | |
|---|---|---|---|---|---|
| 31 | TAFC-Transferrin | TAFC sequesters iron from transferrin bound to iron (TfFe +TfFe2) | Deterministic: reaction proceeds according to Equation 7 | Local levels of TAFC-free and Transferrin Bound to Iron decrease while levels of TAFC Bound to Iron and free transferrin increase. | [130, 69] |
| 32 | Lactoferrin-Transferrin | Lactoferrin sequesters iron from Transferrin Bound to Iron (TfFe + TfFe2) | Deterministic: reaction proceeds according to Equation 7 | Local levels of free lactoferrin and Transferrin Bound to Iron decrease while levels of Lactoferrin Bound to Iron (LTfFe + LTfFe2) and free transferrin increase. | [31, 131] |
| 33 | Macrophage-iron | Necrotic macrophage releases its iron content[2] | Deterministic: whole iron content of the cell is released | Local iron concentration increases | |
| 34 | Aspergillus-Iron | Dead-Aspergillus releases its iron content[2] | Deterministic: whole cell iron content is released | Local iron concentration increases | |
| 35 | TAFC-Iron | TAFC sequesters $Fe^3$ | Deterministic: TAFC sequesters the whole iron content in the voxel or the maximum of its capacity | free TAFC and iron decrease; TAFC bound to iron increases | |
| 36 | Transferrin-Iron | Transferrin sequesters $Fe^3$ | Deterministic: transferrin sequesters the whole iron content in the voxel or the maximum of its capacity | free transferrin and iron decrease; Transferrin Bound to Iron increases | |
| 37 | Lactoferrin-Iron | Lactoferrin sequesters $Fe^3$ | Deterministic: lactoferrin sequesters the whole iron content in the voxel or the maximum of its capacity | free lactoferrin and iron decrease; Transferrin Bound to Iron increases | |

We are using here the macrophage nomenclature used by Gordon, S 2003 [132] where $TGF - \beta$ secreting macrophages are called inactive. This is equivalent to the M2c macrophages referred by Duque, GA & Descoteaux, A 2014 [133].

### S4 Drivers of Fungal Burden

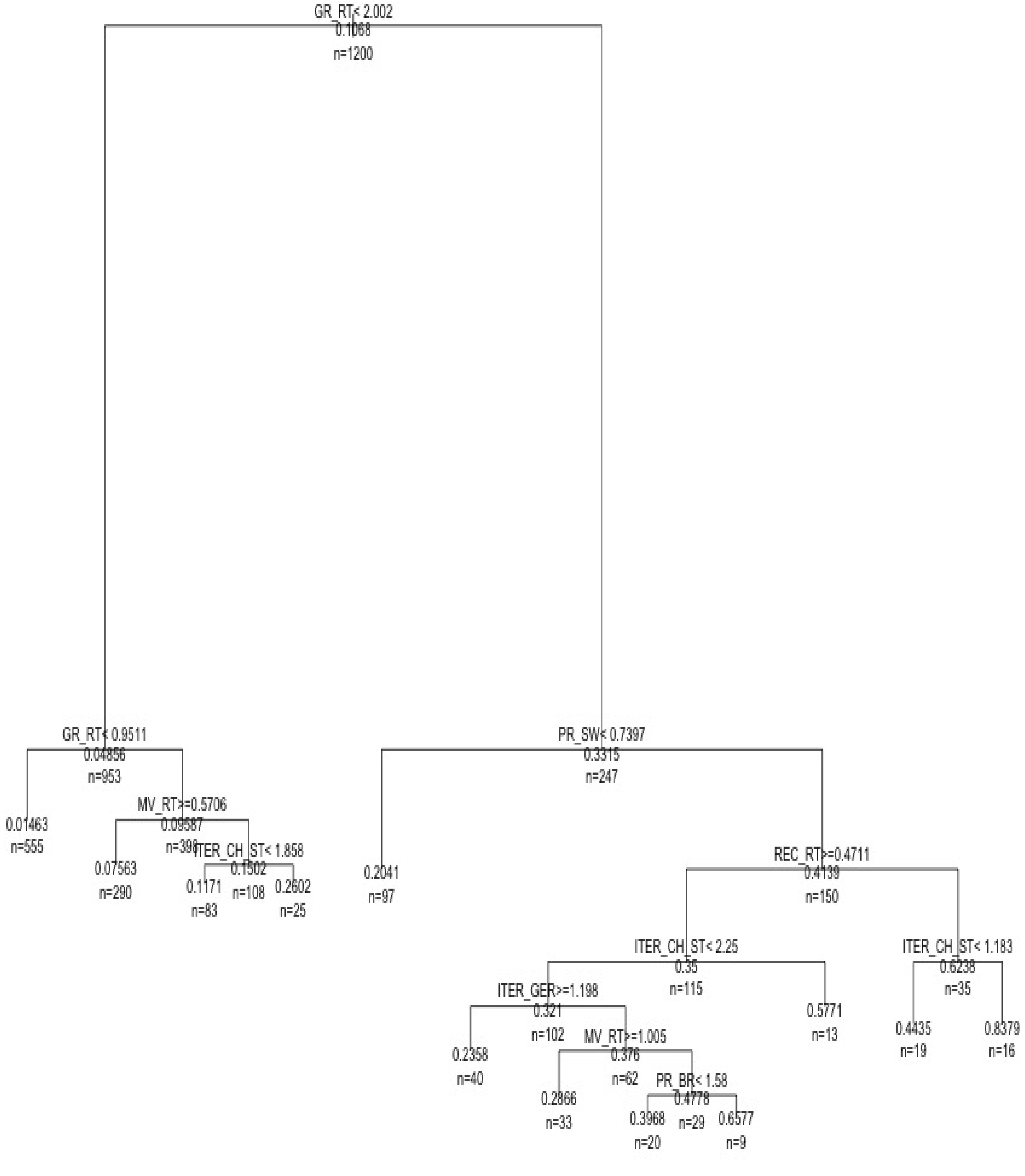


Figure S2: Classification trees of fungal burden. We generated 1200 samples with LHS and ran simulations with immunocompetent hosts, and classified the simulations by the maximum number of hyphal cells measured over 24 hours of simulation. For convenience, all the parameters were normalized by their default values. That is, a value of 2 means that the tree made a partition when that parameter was twice its default value (Table S1). We also normalized the fungal burden by the initial dose inocula (1920 conidia). Notice, however, that the measure of fungal burden is the maximum number of hyphal cells encountered at any time over a 24 hour period. This counts only the conidia that germinate and the hyphae that multiply, excluding ungerminated conidia.

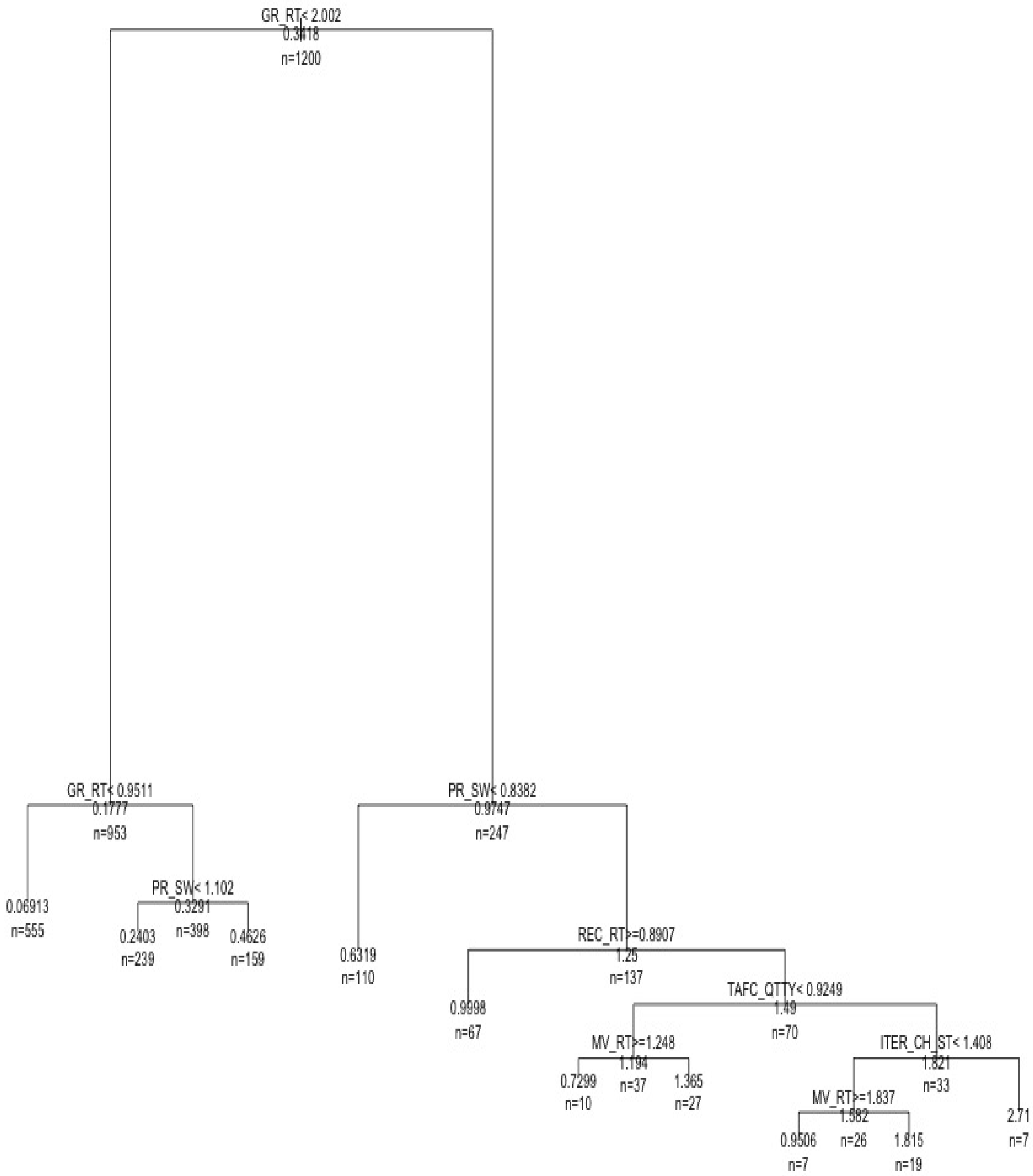


Figure S3: Classification trees of fungal burden. We generated 1200 samples with LHS and ran simulations with neutropenic hosts, and classified the simulations by the maximum number of hyphal cells measured over 24 hours of simulation. For convenience, all the parameters were normalized by their default values. That is, a value of 2 means that the tree made a partition when that parameter was twice its default value (Table S1). We also normalized the fungal burden by the initial dose inocula (1920 conidia). Notice, however, that the measure of fungal burden is the maximum number of hyphal cells encountered at any time over a 24 hour period. This counts only the conidia that germinate and the hyphae that multiply, excluding ungerminated conidia.

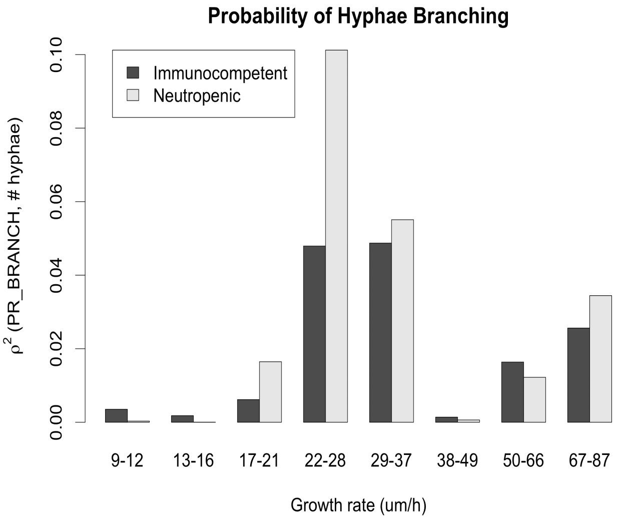


A

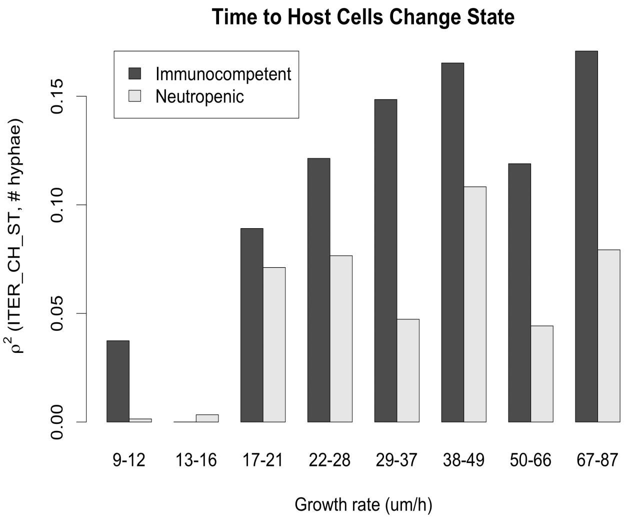


B

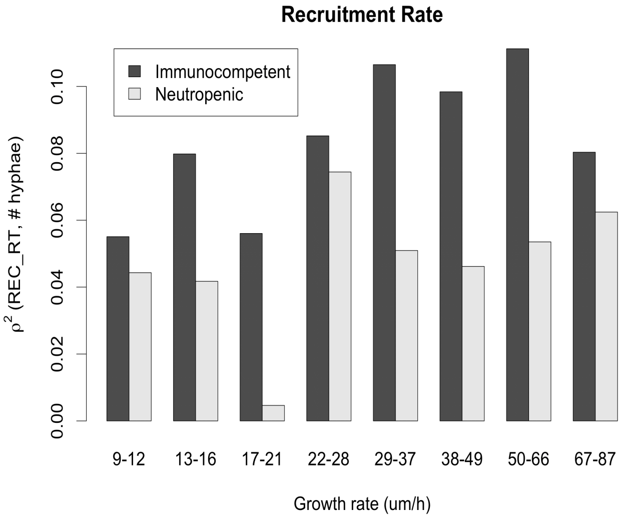


C

Figure S4: Variation of the square of the correlation $r^2$ between the critical model parameters and fungal burden, as the growth rate varies along the x-axis. We started with a dataset of 1200 samples generated with LHS and then partitioned this dataset by growth rate. The x-axis in the figure shows the range of the growth rate in each partition. We then calculate the correlation between parameters and fungal burden in each partition. Fungal burden was the maximum number of hyphal cells measured over the course of 24 hours of simulation. A: variation of $r^2$ between the probability of branching and fungal burden. B: variation of $r^2$ between the iterations for host cells to change state and fungal burden. C: variation of $r^2$ between the leukocyte recruitment rate and fungal burden.
